# Digital Twin Brain simulation and manipulation of a functional brain network underlying mental illness

**DOI:** 10.64898/2026.03.06.710030

**Authors:** Yunman Xia, Songjun Peng, Juergen Dukart, Chao Xie, Shitong Xiang, Spase Petkoski, Zilin Li, Joerg Hipp, Suresh Muthukumaraswamy, Anna Forsyth, Tianye Jia, Nilakshi Vaidya, Tristram Lett, Liyi Qian, Xiao Chang, Yuxiang Dai, Tobias Banaschewski, Gareth J. Barker, Arun L. W. Bokde, Rüdiger Brühl, Sylvane Desrivières, Herta Flor, Penny Gowland, Antoine Grigis, Andreas Heinz, Hervé Lemaître, Frauke Nees, Dimitri Papadopoulos Orfanos, Luise Poustka, Michael N. Smolka, Sarah Hohmann, Henrik Walter, Robert Whelan, Paul Wirsching, Zuo Zhang, Lauren Robinson, Jeanne Winterer, Yuning Zhang, Hedi Kebir, Ulrike Schmidt, Julia Sinclair, Yuchen Liu, Jiexiang Wang, Fei Dai, Longbin Zeng, Yubo Hou, Huarui Wang, Leijun Ye, Chunhe Li, Qibao Zheng, Andre Marquand, Changsong Zhou, Viktor Jirsa, Jianfeng Feng, Wenlian Lu, Gunter Schumann, IMAGEN Consortium, STRATIFY Consortium, DTB Consortium, environMENTAL Consortium

**Author notes:** These authors contributed equally to this work (Yunman Xia, Songjun Peng). Corresponding author: Gunter Schumann, and.

## Abstract

Linking synaptic-level perturbations to distributed brain-network dynamics remains a central challenge for understanding and treating mental illness. Although recent whole-brain models can reproduce individual brain activity patterns, they largely function as descriptive simulators rather than mechanistic, intervention-capable systems. Here we present an intervention-capable digital twin of the human brain, integrating individual neuroanatomy and task-evoked dynamics within a neuronal-scale framework. Individualised digital twin brains recapitulate a participant-specific compact cortico-subcortical network phenotype that captures transdiagnostic psychopathology across population and clinical cohorts. In silico modulation of excitatory and inhibitory synaptic conductance produces bidirectional, heterogeneous network responses across individuals. Population-scale simulations stratify individuals and predict longitudinal symptom trajectories from DTB-derived response profiles. Independent pharmacological functional MRI data further validate the predicted baseline-dependent network responses in vivo. Together, these findings establish digital brain models as experimental platforms for mechanistic perturbation, behavioural prediction and stratification, providing a foundation for precision neuroscience and psychiatry.

## Main

As human neuroimaging has reached individual-level precision^1,2^, a central challenge has emerged: how to transform rich multimodal measurements of brain structure and function into mechanistic models that can be systematically perturbed to generate causal and predictive insights into highly prevalent mental illness^3,4^. Digital twin models of the human brain offer a powerful framework for addressing this challenge by integrating neuroanatomy, physiology, and large-scale functional dynamics within a unified computational system^5–8^. Recent advances have demonstrated that Digital Twin Brain (DTB) framework can approximate large-scale neuronal dynamics using the biologically constrained spiking neural networks grounded in individual anatomy and empirical BOLD signals^9,10^, establishing the feasibility of personalised whole-brain simulation. However, existing digital twin brains primarily function as descriptive simulators rather than enabling systematic mechanistic perturbation and prediction.

This limitation is especially salient in psychiatry. Mental disorders are characterised by distributed network alterations^11^ and heterogeneous treatment response^12–15^. Neuroimaging studies have identified reproducible functional connectivity abnormalities across disorders, often implicating common circuits involved in reward processing, cognitive control, and salience attribution^15–22^. Yet without the ability to perform controlled perturbations, these findings remain difficult to translate into mechanistic insight or predictive models of intervention effects^13,14^. At the same time, most psychiatric treatments act on specific molecular targets - such as glutamatergic or GABAergic neurotransmission - whose effects propagate across multiple spatial scales, from synapses to distributed functional brain networks^23–27^. Bridging these levels of description remains a critical gap.

A promising strategy is to focus on behaviourally grounded, transdiagnostic brain networks that provide low-dimensional interfaces between cognition, symptoms, and neurobiology^18–20,28^. Converging evidence suggests that shared dimensions of psychopathology are underpinned by common neural circuit disruptions that cut across traditional diagnostic boundaries^17–20,28–30^. The neuropsychopathology (NP) factor, derived from task-based functional connectivity, captures shared continuous variance across internalising and externalising symptoms in the population cohort^28^. As such, it provides a principled target for testing whether digital twin brains can move beyond simulation toward mechanistic manipulation of behaviourally relevant circuits.

In parallel, a large body of neurochemical evidence implicates disruptions in cortical excitation–inhibition (E/I) balance in the pathophysiology of mood and addiction disorders. Magnetic resonance spectroscopy and postmortem studies report reductions in both glutamatergic and GABAergic signalling across cortex^31–36^, while pharmacological agents targeting these systems - including ketamine and benzodiazepines - can induce rapid changes in symptoms and functional connectivity^23,26,37–41^. Despite this, the relationship between neurotransmitter-level perturbations and large-scale functional network reconfiguration remains poorly understood. Empirical pharmaco-fMRI can quantify individual-level effects of neurochemical interventions, but offers limited leverage to mechanistically disentangle or systematically manipulate the underlying synaptic processes. Existing whole-brain models, in turn, have not yet been configured to support controlled, neurotransmitter-level perturbations to task-state network reconfiguration at the individual level^8,42–44^.

Here we build on the DTB framework^9,10^ to address this unmet need. We tested whether a neuronal-scale digital twin can reproduce a behaviourally relevant transdiagnostic brain network (NP factor) and mechanistically manipulate it through abstract synaptic gain control parameters inspired by E/I mechanisms. We further examined whether such in silico manipulations yield predictions that generalise across tasks and populations, align with empirical pharmacological effects, and forecast longitudinal symptom trajectories. Together, this work marks a step change in digital brain modelling from passive emulation of neural activity to intervention-capable, mechanistically closed-loop systems that integrate simulation, perturbation, and prediction.

### A compact cortico–subcortical network phenotype of psychopathology

We first delineated the NP factor, a shared functional connectivity phenotype that captures psychopathology across symptom domains and cohorts. As in our previous work^28^, conducted in 14 year-old adolescents of the IMAGEN study, we applied connectome-based predictive modeling to the task-based functional connectomes derived from the monetary incentive delay and stop-signal task in 19 year old young adults of IMAGEN follow-up 2 (n=1,050, age 18.4±0.7 years, Table S1). Four of six task conditions (stop success and failure in SST; reward anticipation and positive feedback in MID) significantly predicted a range of externalising and internalising symptoms at age 19 (Fig.1a; Table S2-3). Across these predictive task conditions, we identified two transdiagnostic functional connectivity (FC) profiles: one positive profile of 34 edges where stronger connectivity was associated with higher symptom burden across domains, and one negative profile of 29 edges where weaker connectivity was associated with higher burden. These edges were concentrated within and between default mode, frontoparietal, somatomotor–frontal, limbic, and cerebellum systems (Fig.1b). To quantify these functional connectivity profiles, we computed positive and negative NP scores for each individual by summing the strength of edges in each profile.

**Figure 1.**
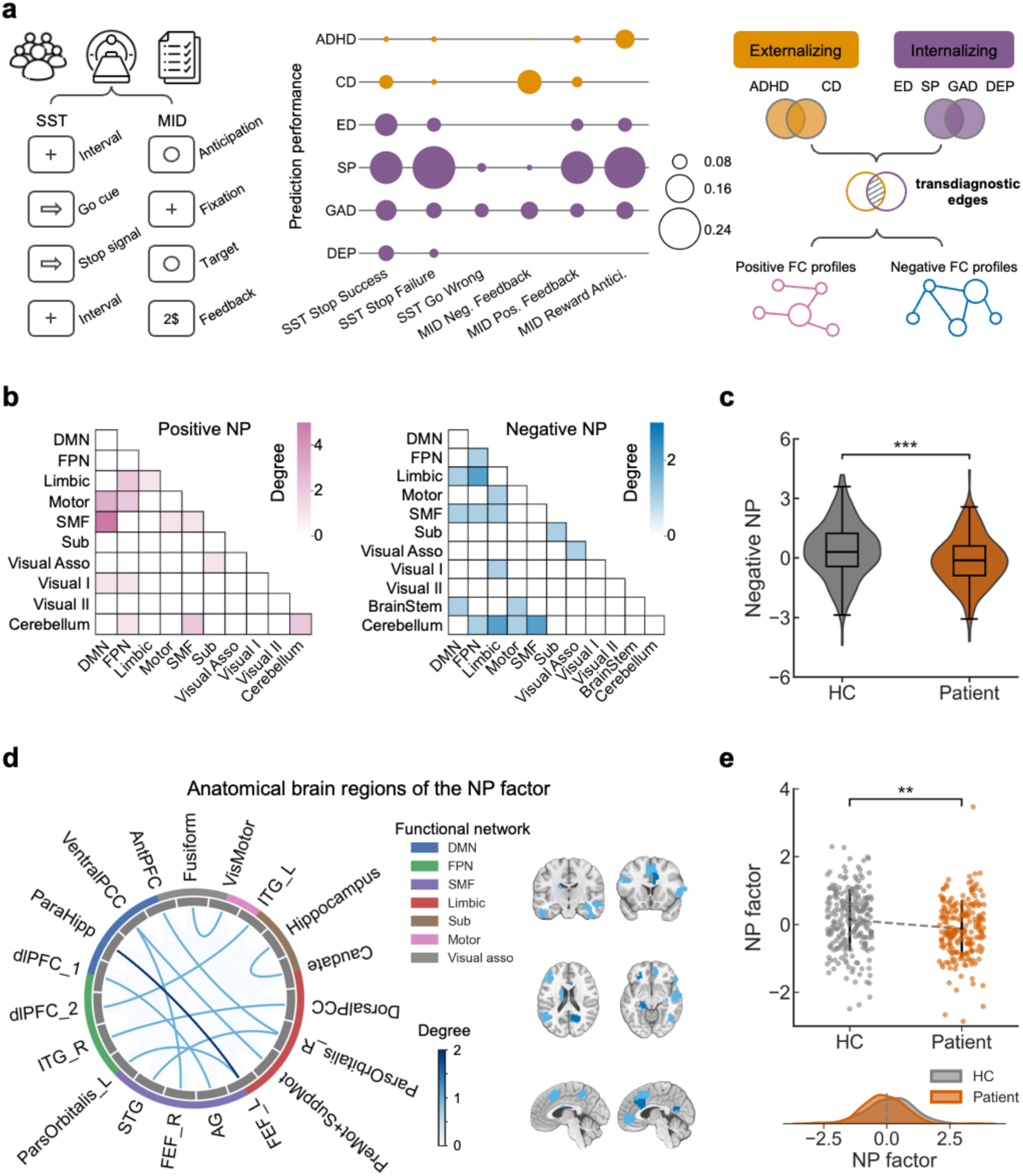
Identification of a compact functional network phenotype of transdiagnostic psychopathology. **a,** Task-based fMRI data and behavioural symptom scores from the IMAGEN cohort (n = 1,050) were analysed using connectome-based predictive modelling (CPM) to identify task-evoked functional connectivity (FC) profiles associated with transdiagnostic psychopathology. Predictive performances of task-specific FC derived from the Stop-Signal Task (SST) and Monetary Incentive Delay (MID) were shown in a bubble plot, where bubble size reflects Spearman’s correlation between predicted and empirical symptom scores. The mean *P* values across 1,000 repetitions were corrected for multiple comparisons across 36 models using false discovery rate (FDR) correction, corresponding to a minimum significant effect size of Spearman’s *ρ* = 0.07. Task-specific functional connectivity (FC) matrices were then selected if they showed significant prediction performance for at least three behavioural symptom domains. SST conditions included stop-success, stop-failure, and go-wrong; MID conditions included positive feedback, negative feedback, and reward anticipation. Behavioural measures comprised two externalising domains (attention-deficit/hyperactivity disorder, ADHD, and conduct disorder, CD) and four internalising domains (eating disorder, ED, specific phobia, SP, generalized anxiety disorder, GAD, and depression, DEP). A transdiagnostic FC profile was identified that significantly predicted both externalising and internalising symptoms. These transdiagnostic associated edges were divided into positive (pink) or negative (blue) FC profiles based on the direction of their associations with behavioural symptoms**. b,** Heatmaps show the canonical functional network affiliations of positive and negative transdiagnostic FC profiles. Positively predictive connections were primarily distributed across the default mode network (DMN), frontoparietal network (FPN), somatomotor–frontal (SMF) network, and cerebellum, whereas negatively predictive connections were concentrated in limbic, DMN, FPN, SMF, and cerebellar systems. **c,** Negative NP scores (summed strength of the negative FC profile) were compared between healthy controls (HC, n = 225) and patients (n = 209) from the STRATIFY cohort. Group difference was assessed using an independent-samples *t* test with Bonferroni correction (*t* = 3.94, *P*_corrected_ = 0.0003). Values represent residuals after regressing out sex, scanning site, and head motion (mean framewise displacement). Violin plots show kernel density estimates with embedded boxplots indicating the median and interquartile range (IQR); whiskers extend to 1.5× IQR. **d,** After excluding cerebellar and brainstem connections, the negative FC profile was reduced to a compact 12-edge NP circuit comprising cortico–subcortical connections. Edges in the circos plot represent connection strength and negative associations with psychopathology; corresponding brain regions are shown on cortical and subcortical renderings. **e,** Individual summed 12-edge-NP factor scores were compared between HC and patients from the STRATIFY cohort. Group difference was confirmed using an independent-samples *t* test with Bonferroni correction (*t* = 3.27, *P*_corrected_ = 0.0036). Scores represent residuals after regressing out sex, scanning site, and head motion. Each dot denotes one participant; black markers and vertical lines indicate group means and standard deviations, respectively. Kernel density estimates illustrate the distributional shift in NP factor scores between groups. **, *P*_corrected_ < 0.01; ***, *P*_corrected_ < 0.001.

To test whether the NP scores generalise to clinical populations, we re-computed them from task-state FC in an independent case–control cohort (STRATIFY, n=434, age 21.8±2.0 years). Only the negative NP score showed robust case-control differences, with lower scores in patients than in healthy controls (Fig.1c; Fig.S1). Thus, the negative NP profile defines a shared network phenotype of transdiagnostic psychopathology.

To characterise the neurobiological context of this negative NP circuit, we examined whether negative NP-related regions exhibit a distinctive neurochemical signature in empirical PET receptor maps. Using published PET-derived receptor density maps^45^, we found that NP-related regions showed higher receptor-map values for glutamatergic, serotonergic, cholinergic, and cannabinoid systems than NP-unrelated regions (Fig.S2, Table S4, all *P*_corrected_ < 0.05). These receptor systems are primarily involved in regulating cortical excitatory drive and neuromodulatory control of network dynamics, and are known to influence E/I balance at the circuit level through modulation of glutamatergic transmission^46,47^, GABA interneuron recruitment^48–50^, recurrent coupling^51^, and gain control^52^ mechanisms.

To enable computationally tractable digital twin brain (DTB) modeling, we restricted the negative NP profile to cortico-subcortical edges, excluding the cerebellum and brainstem. These regions exhibit distinct cytoarchitecture and neuronal dynamics^53^ and are not currently parameterized in our DTB framework (see Methods for details). Finally, we identified a compact set of 12 task-state functional connections linking regions spanning prefrontal control, default-mode, sensorimotor, and limbic systems (Fig.1d; Table S5). The summed strength of these 12 edges remained significantly reduced in patients relative to controls (Fig.1e; Fig.S1), and served as the final NP factor and target for subsequent DTB simulations and in-silico E/I modulations.

### Individual digital twin brains reveal a bidirectionally E/I-tunable NP factor

We next asked whether this NP factor can be reconstructed and mechanistically manipulated in individualised, biologically grounded models. To this end, we applied the previously established DTB framework^9,10^ to generate participant-specific task-state models (Methods; Fig.2a; Fig.S3). Following a calibration experiment across neuronal resolutions (Fig.S4; Table S8; more details in Methods and Supplementary Results), we implemented DTBs at three complementary scales: billion-neuron voxel-wise models to establish biological fidelity and simulation feasibility, 100-million-neuron models for controlled perturbational exploration, and computationally tractable regional models for population-level statistical inference. Across all scales, models were constructed using a common pipeline. The individual voxel-wise grey and white matter anatomy were first transformed into a multi-population neuronal network. The activity of each neuron in this network was modelled using the leaky integrate-and-fire (LIF) model^54^, equipped with excitatory (AMPA-mediated) and inhibitory (GABA-A-mediated) synapses. We then estimated hyperparameters by assimilating empirical BOLD signals in task-engaged regions (Methods; Table S6) to mimic external current inputs driving the model from rest to task-states^9,10^. Finally, the voxel-wise neuronal activity (mean firing rate of neuronal population) was converted into BOLD signals using the Balloon-Windkessel model^55^.

**Figure 2.**
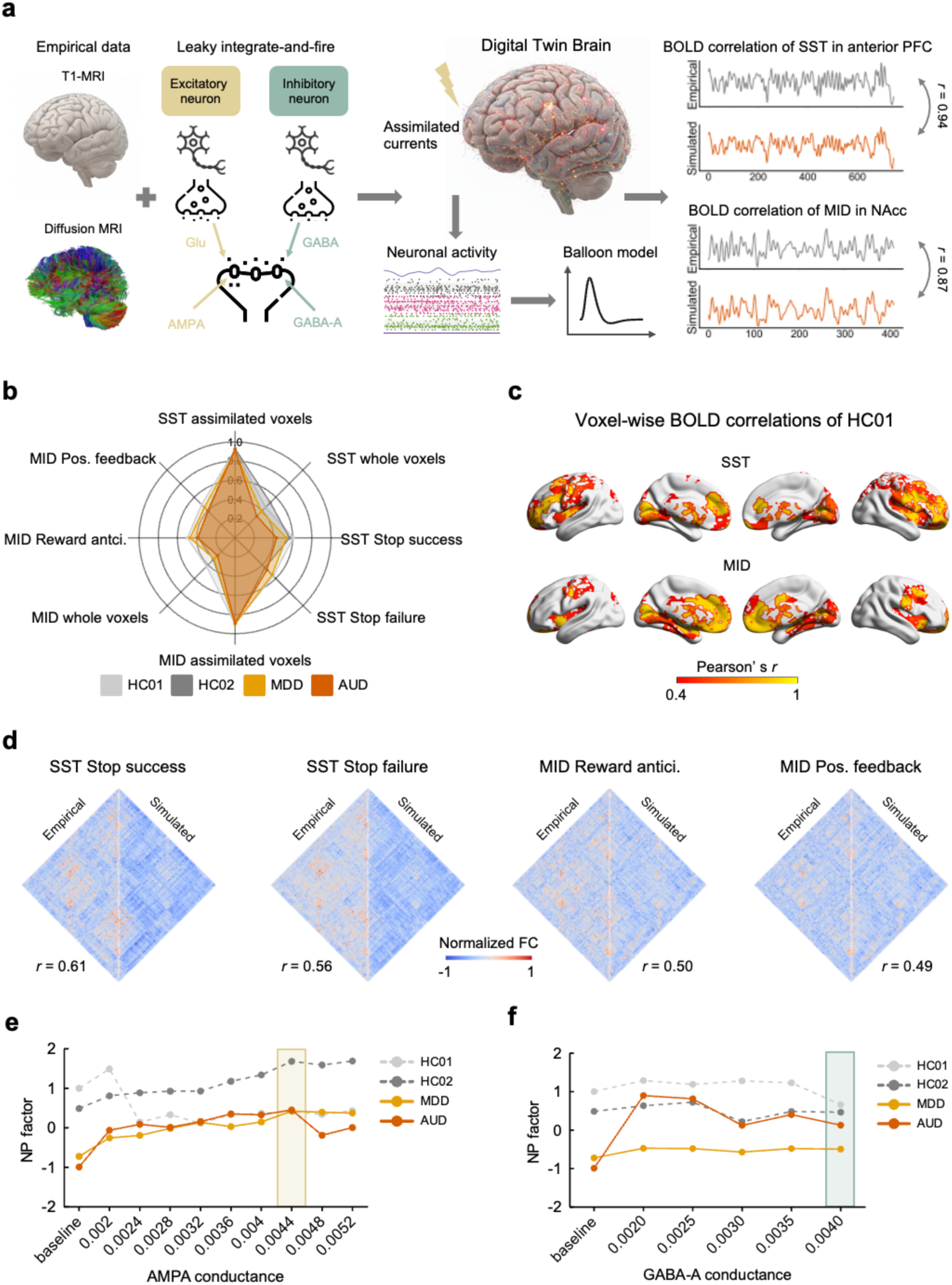
Individualised digital twin brains (DTBs) reveal excitation/inhibition-sensitive NP modulation. **a,** Computational workflow for constructing individualised DTBs from multi-modal participant-specific MRI data. DTB model received assimilation hyperparameters as external current inputs and generated simulated neuronal firing rates, which were converted to BOLD signals using Balloon model. The simulation performance was assessed by Pearson’s *r* between empirical and simulated BOLD signals. **b,** Radar plots showing simulation performances of 1-billion-neuron models for two patients (one MDD; one AUD) and two healthy controls (HC01; HC02). For each task, both the mean voxel-wise Pearson’s correlations in assimilated voxels (assimilated) and the mean correlations across the whole voxels (whole). Task-specific FC matrices included stop-success and stop-failure for the SST, and reward anticipation and positive feedback for the MID task. **c,** Voxel-wise Pearson’s correlations between empirical and simulated BOLD signals from the 1-billion-neuron DTB model for the MID and SST tasks of HC01. Bright yellow regions indicate higher similarity. **d,** Empirical and simulated FC matrices across four task conditions of SST and MID from the 1-billion-neuron DTB model of the HC01. **e–f,** In silico E/I modulation sweeps. Curves depict NP factor responses to systematic changes in AMPA (**e**) and GABA-A (**f**) synaptic conductance. For AMPA modulation, NP factor scores increased in patients and HC02 but decreased in HC01. For GABA-A modulation (with AMPA = 0.0044, the parameter yielding maximal NP factor improvement), the NP factor scores increased in two patients, while slightly decreased in HCs. The shaded areas indicate optimal E/I parameters for future perturbations (AMPA = 0.0044; GABA-A = 0.0040). **, *P_corrected_* < 0.01.

We first selected four participants as illustrative exemplars according to the behavioural symptoms and negative NP scores, including participants with major depressive disorder (MDD), alcohol use disorder (AUD), and healthy controls (HC01 and HC02) (demographics information in Table S7, selection criteria in Methods), to demonstrate the feasibility and mechanistic properties of neuronal-scale digital twin simulations. These examples are presented for illustration only; all statistical inferences are based on group-level analyses that account for site and sex effects (see Supplementary Results). Using a 1-billion-neuron scale (Fig.S4; Table S8), their individualised DTBs accurately reproduced task-evoked dynamics in MID and SST. Simulations achieved high voxel-wise correlations with empirical BOLD in assimilated regions (mean *r*=0.9, in-sample correlations). Across the whole brain, mean correlations of four individuals (mean *r*=0.30-0.44) were clearly above a reference simulation obtained by a single random parameter shuffle (mean *r*=0) (Fig.2b-c; Fig.S5-6; Table S9). These DTBs also produced whole-brain functional connectivity patterns that closely matched empirical connectomes in both tasks (Fig.2d; Table S9).

Based on the simulated baseline NP configuration, we next examined whether NP circuit dynamics could be systematically manipulated within the DTB framework. In recurrent cortical networks, E/I balance emerges from the interaction of inhibitory interneuron recruitment^56,57^, recurrent coupling^51,57^, and synaptic gain^58^, and is governed by circuit dynamics rather than receptor density alone. Consistent with this systems-level perspective, we treated both AMPA- and GABA-A-mediated synaptic conductance as global dynamical control parameters, enabling controlled perturbation of NP network activity within individualised digital twins. Notably, publicly available PET datasets did not show preferential expression of GABA receptor systems in NP-related regions, reinforcing the interpretation that inhibitory modulation in the model reflects circuit-level dynamical control rather than regionally localized receptor enrichment.

Accordingly, we varied the synaptic conductance parameters for AMPA and GABA-A receptors in the 100-million-neuron DTBs of the same illustrative participants. Despite the reduced resolution, DTBs preserved high fidelity in assimilated regions and acceptable whole-brain correspondence (Table S10). Moreover, 100-million-neuron DTBs retained sufficient subject-specific NP-factor phenotypes such that each individual’s simulated NP factor best matched their own empirical value, with significantly lower mean-squared error for within-subject matches than any cross-subject pairing (permutation tests across 1,575 simulated–empirical pairings, Supplementary Results; Fig.S7), therefore supporting its use for controlled perturbational experiments. Using these established DTB models, we performed systematic perturbations by upregulating synaptic conductance for AMPA (0.0020-0.0052) and GABA-A receptors (0.0015-0.0040) across a predefined parameter grid, guided by evidence for reduced glutamatergic and GABAergic function in depression^24,25^ and addiction^27,34^. Parameter ranges were constrained by biologically plausible neuronal firing rates^59^, synchronization^60^, and simulated BOLD activity. During GABA-A manipulations, AMPA conductance was fixed at the value that maximized NP factor enhancement in prior sweeps, allowing controlled assessment of E/I balance and avoiding excessive inhibition (see more details in Methods).

Virtual modulation of AMPA- and GABA-A-mediated synaptic conductance produced bidirectional, heterogeneous changes in NP connectivity across illustrative participants. Specifically, in two patient brain models, both excitatory and inhibitory upregulation increased NP-factor strength, and in illustrative healthy control models, the same perturbations yielded divergent responses, with modest increases in one individual and clear decreases in the other (Fig.2e-f; Tables S11-12). These examples demonstrate that identical synaptic perturbations can lead to bidirectional changes in NP-factor strength across individuals.

To test whether this bidirectional response generalises across cognitive contexts, we repeated the same E/I perturbations in an alternative task (the Emotional Face Task) in the same participants. Comparable bidirectional modulation effects were observed (Supplementary Results; Fig.S8; Table S13), indicating that the E/I-sensitive response properties of the NP circuit are not task-specific, but generalise across distinct cognitive contexts.

### Population-scale in silico perturbations attenuate NP abnormalities and stratify individuals

The single-subject analyses suggest that in silico E/I perturbation can shift NP network toward the control distribution observed in STRATIFY (reflected by increased NP-factor strength), although not uniformly across individuals. To further explore whether these effects generalise across the cohort and relate to behavioural differences, we performed population-scale simulations, which allow us to link individual susceptibility to excitatory and inhibitory synaptic perturbations with behavioural symptoms. To enable large-cohort simulations, we developed regional DTBs that convert regional MRI-derived structural features into spiking neural networks comprising 3 million neurons (identified from calibration experiment, Fig.S4, Table S14), with neuron counts allocated proportionally to regional grey matter volume to preserve anatomical scaling while maintaining computational tractability (Methods). While voxel-wise DTBs establish mechanistic feasibility at high-fidelity resolution, these regional DTBs are used to test whether these principles scale to population-level inference.

We constructed individualised DTBs for 72 patients with MDD, 59 with AUD and 69 healthy controls from STRATIFY, as well as 90 individuals with high externalising and internalising symptoms from IMAGEN (Table S15). These participants from STRATIFY were included because they possessed complete, high-quality multimodal imaging (T1-weighted, DTI, fMRI) and behavioural data. These DTB models reproduced individual task-evoked BOLD signals with moderate-to-high accuracy during in-sample fitting phase (Fig.3a), generated NP factors that closely matched empirical values at the group level (Fig.S9), and mirrored case–control group differences observed in vivo (Fig.3b; Fig.S9). As an additional validation beyond NP circuit, simulated whole-brain FC also predicted task-related behavioural performances and, on average, outperformed empirical FC (Supplementary Results; Fig.S10; Table S16), suggesting that DTBs distill behaviourally relevant variance that is only partially captured by conventional functional connectivity.

**Figure 3.**
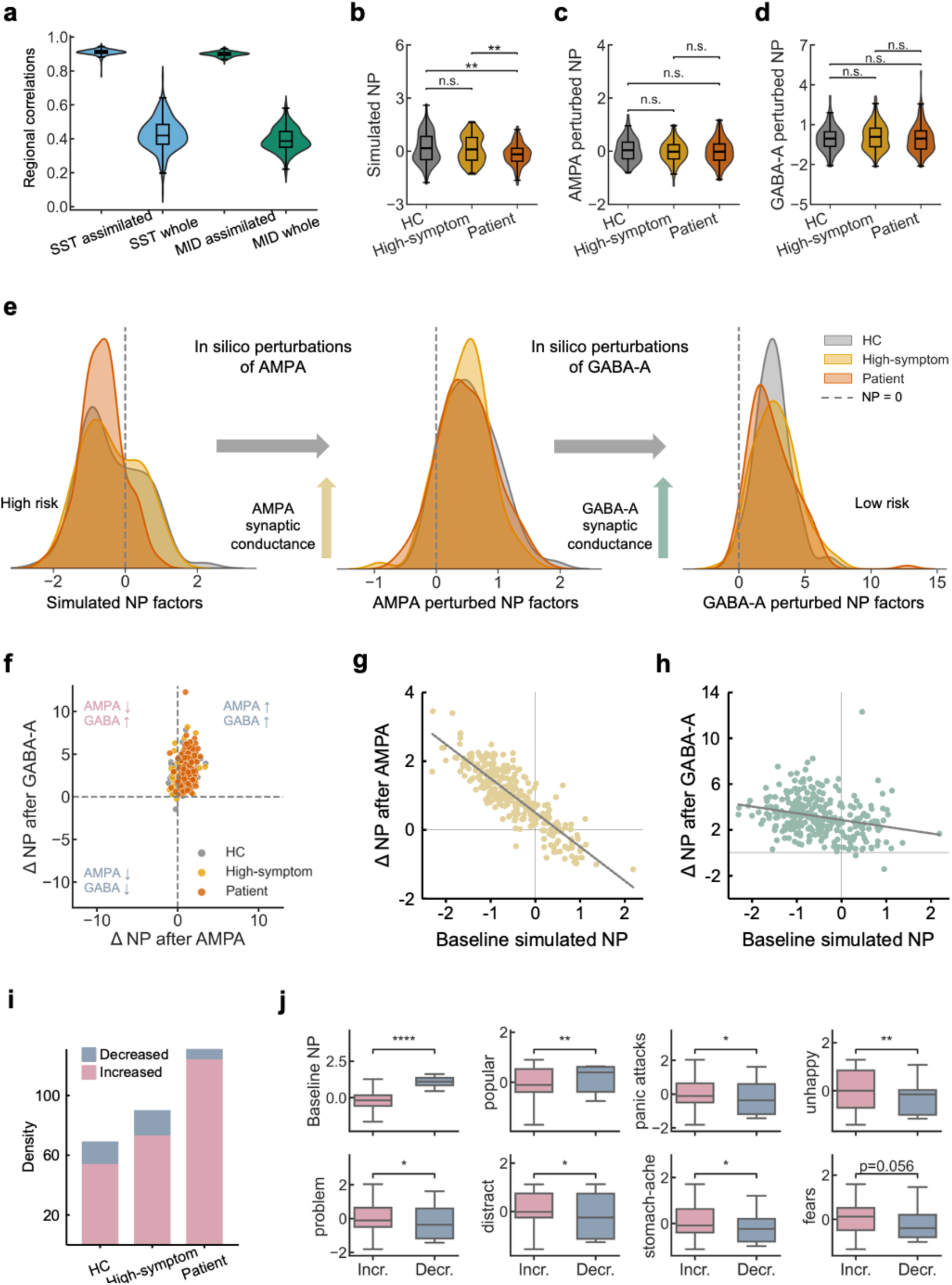
Population-scale in silico perturbations and mechanistically grounded stratification. **a,** Regional mean Pearson’s correlations between empirical and simulated BOLD signals during SST and MID tasks from 3-million-neuron DTB models for 290 participants. For each task, Violin plots show both the mean correlations within assimilated task-related regions (assimilated) and the mean correlations across all regions (whole). The boxplots indicating the median and IQR; whiskers extend to 1.5× IQR. **b-d,** Simulated NP factor scores under baseline (**b**), AMPA-mediated (**c**), and GABA-A-mediated (**d**) virtual perturbations in healthy controls (HC, n = 69), high-symptom individuals (n = 90), and patients (n = 131). Group differences were assessed using ANOVA and post-doc tests with Bonferroni correction for multiple comparisons (n.s., not significant; **, *P_corrected_* < 0.01). Values are residuals after regressing out sex, site, and head motion. Violin plots show kernel density estimates with embedded boxplots indicating the mean and IQR; whiskers extend to 1.5× IQR. **e,** Kernel density distributions of NP factor scores across healthy controls (HC, grey), high-symptom individuals (yellow), and patients (orange) under three simulation conditions: (left) baseline, (middle) AMPA modulation, and (right) GABA-A modulation. The x-axis represents the simulated NP factor scores, and shaded areas denote the distribution density for each group. Across the three panels, the degree of overlap between group distributions and the relative shift of scores from negative to positive values correspond to the transition from baseline to specific receptor manipulations. **f,** Individual changes in NP factor scores following the AMPA and GABA-A perturbations. Dashed lines indicate zero change. Each dot represents an individual subject. Individuals with ΔNP>0 (perturbed-baseline) under both perturbations were classified as the increased-response group (n = 251, pink); others formed the decreased-response group (n = 39, blue). **g-h,** Baseline NP factor scores were inversely correlated with AMPA-(**g**, Pearson’s *r* = -0.86, *P* < 0.0001) and GABA-A-(**h**, Pearson’s *r* = -0.25, *P* < 0.0001) induced changes, with a permutation test of 10,000 times. **i,** Distribution of response types across groups (Chi-square test: *χ²* = 14.05, *P* = 0.007). Increased-response proportions: patients, 124/131 (94.66%); high-symptom, 73/90 (81.11%); HC, 54/69 (78.26%). **j,** The increased-response group exhibited lower baseline NP factors, and higher symptom burden, including panic attacks, negative affect and reduced social support, than the decreased-response group (Mann–Whitney U tests, controlling for sex, site, and head motion; *, *P*<0.05; **, *P*<0.01; ****, *P*<0.0001). Boxplots represent the median and IQR; whiskers extend to 1.5× IQR.

Motivated by these converging validations, we then applied the same AMPA and GABA-A neuromodulation schemes used in the 100-million-neuron DTBs to all 290 participants. At the group level, both excitatory and inhibitory upregulation shifted NP factors in patients toward values observed in controls, in other words, after either AMPA or GABA-A modulation, previously significant differences between patients, high-symptom individuals and healthy controls were no longer significant (Fig.3c-d). Within each clinical group, NP factors increased on average following both AMPA and GABA-A manipulations (Fig.3e; Fig. S11).

Despite clear group-level increases, individual responses were heterogeneous and often bidirectional (Fig.3f). Following AMPA modulation, both patients and controls showed increased NP factor on group average, but the proportion of “increasers” (ΔNP > 0) was substantially higher among patients than controls (Table S17). Baseline simulated NP factor was strongly and inversely correlated with AMPA-induced change (Fig.3g), such that individuals with lower baseline connectivity, typically patients, exhibited the largest gains, whereas those with high baseline NP showed modest increases or decreases. GABA-A modulation produced a similar, though weaker, negative association (Fig.3h). Importantly, these baseline–ΔNP couplings significantly exceeded a null distribution generated by permuting ΔNP values across individuals (10,000 permutations, two-sided *P*<0.0001; see Methods), indicating that the relationship reflects structured baseline-dependent dynamics rather than statistical regression to the mean.

To facilitate clinical interpretation of this response heterogeneity, we descriptively stratified individuals based on the direction of their joint responses to AMPA and GABA-A perturbations. Participants whose NP factors increased under both manipulations were classified as “increased responders”, whereas those with at least one decrease were classified as “decreased responders”. This stratification was used for visualization and group-level comparison only; the underlying baseline–ΔNP relationships were treated as continuous throughout statistical analyses. Increased responders were markedly more prevalent among patients than among high-symptom individuals or healthy controls (Fig.3i) and did not differ systematically by site or sex (Tables S18–19; Supplementary Results). Clinically, the increased responders, characterised by lower simulated baseline NP factors, showed greater symptom burden, including more panic attacks and negative affect, and reduced social support, compared with the decreased responders after controlling for sex, site, and head motion (Fig.3j; Table S20).

Together, these findings indicate that DTB-derived virtual responses offer an individualised, mechanistically interpretable stratification, whereby different responses in NP circuit map onto meaningful differences in clinical and behavioural profiles. A critical question, however, is whether these virtual baseline-dependent, bidirectional response structures also emerge under real pharmacological perturbations in vivo.

### Pharmacological E/I perturbation in vivo mirrors digital twin predictions of NP modulations

To address this, we analysed pharmacological fMRI data from 27 healthy male participants (27.3±6.2 years), who performed the MID task following ketamine (0.25 mg/kg), midazolam (0.03 mg/kg), or placebo administration in a randomized, single-blinded, three-way crossover design (Fig.4a). Ketamine, an NMDA receptor antagonist, reduces inhibitory interneuron activity and indirectly enhances AMPA-mediated excitatory transmission^23,61^, whereas midazolam, a GABA-A receptor positive allosteric modulator, increases inhibitory synaptic conductance and dampens network excitability^62,63^. Because pharmacological data were only available for the MID task-fMRI, we focused on a MID-specific NP connectivity, defined as the summed strength of the six MID edges that contributed to the NP circuit. Placebo served as the empirical analogue of the DTB baseline simulation, whereas ketamine and midazolam provided in vivo perturbations with predominant effects on excitatory-leaning and inhibitory neurotransmission, respectively. We tested whether these perturbations reproduce the baseline-dependent response structures predicted by AMPA and GABA-A modulations in silico.

**Figure 4.**
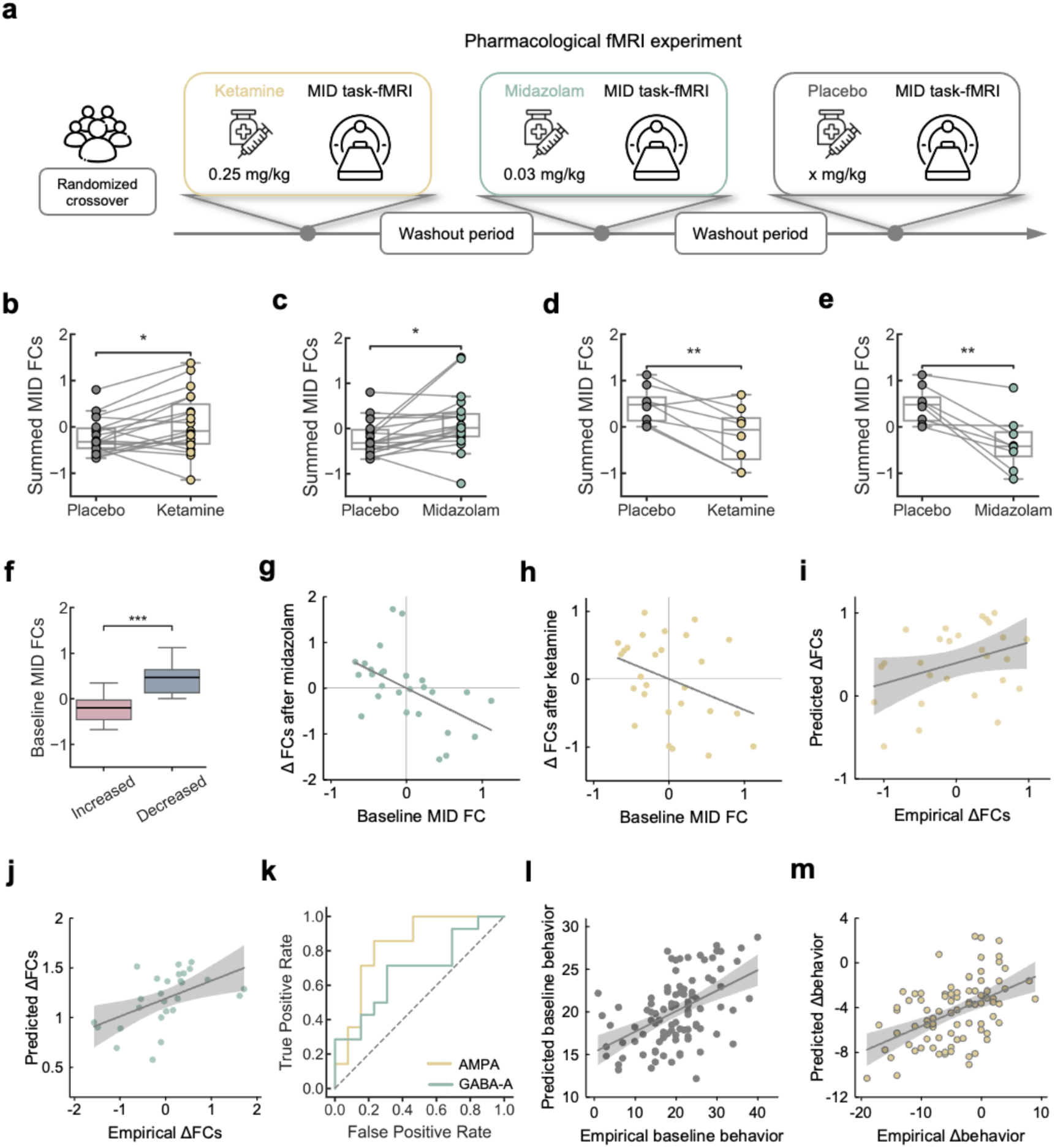
Pharmacological fMRI validation and prospective symptom prediction. **a,** Pharmacological fMRI experiment in 27 male healthy volunteers using a three-way randomized, controlled crossover design. **b-e,** Individual- and group-level changes in summed NP-related MID functional connectivity (summed MID FCs) following ketamine and midazolam administration in subgroup 1 (**b**, **c**, n = 19) and subgroup 2 (**d**, **e**, n = 8). Each dot represents one individual and is colored by drug condition: placebo (grey), ketamine (gold), and midazolam (teal). Statistical significance was assessed using paired *t*-tests or Wilcoxon signed-rank tests, as appropriate. Values represent residuals after regressing out head motion. **f,** Subgroup 1, exhibiting drug-induced increased responses, showed lower baseline summed MID FCs under placebo compared with subgroup 2 (Mann–Whitney U tests, controlling for head motion; ***, *P*<0.001). **g-h,** Baseline summed MID FC was inversely correlated with midazolam-induced change (**g**, Pearson’s *r* = -0.52, 10,000 permutations, *P* = 0.0056), and showed a similar trend for ketamine-induced change (**h**, Pearson’s *r* = -0.35, 10,000 permutations, *P* = 0.069). **i-j,** Prediction of empirical drug-induced FC changes from baseline summed MID FC using linear mappings between DTB-simulated E/I perturbations and simulated baseline MID FC strength. Scatter plots show predicted versus observed FC changes induced by ketamine (**i**, Pearson’s *r* = 0.35, 1,000 permutations, *P* = 0.039) and midazolam (**j**, Pearson’s *r* = 0.52, 1,000 permutations, *P* = 0.001). Each dot represents one individual; solid lines indicate linear regression fits with 95% confidence intervals. **k,** Prediction of drug-induced response direction (ΔNP>0 or <0) from baseline summed MID FCs using DTB-derived models (AMPA model AUC = 0.69; GABA-A model AUC = 0.83). **l,** Linear association between empirical NP factor scores and behavioural symptom severity at age 19 (n = 100, *r²* = 0.24, *F* = 2.3, *P* = 0.014). Each dot represents one individual; solid lines indicate linear regression fits with 95% confidence intervals. **m,** Prospective prediction of behavioural symptom changes over 4 years. Scatter plot shows predicted versus empirical behavioural changes from age 19 to 23. The prediction model explained a significant proportion of variance (n = 85, *r²* = 0.23, *F* = 4.80, permutation test of 5,000 times, *P* = 0.0006). Each dot represents one individual; solid line indicates the linear fit with 95% confidence intervals. All boxplots show the median and interquartile range (IQR); whiskers extend to 1.5× IQR. *, *P* < 0.05; **, *P* < 0.01; ***, *P* < 0.001.

At the group level, neither ketamine nor midazolam induced a significant change in summed NP-related MID connectivity relative to placebo (Fig.S12), suggesting that conventional group-mean analyses would miss potential relationship between brain network configuration and pharmacological response. However, individual-level analyses revealed a bidirectional, baseline-dependent pattern closely resembling DTB simulations. Unsupervised k-means clustering of combined ketamine- and midazolam-induced FC changes identified two subgroups (Fig.S12). In Group 1 (n=19), both ketamine and midazolam significantly increased NP-related connectivity after adjusting for head motion effects (Fig.4b-c), whereas in Group 2 (n=8), both drugs significantly decreased it (Fig. 4d-e). Baseline NP-related connectivity under placebo was significantly lower in the increased-response group than in the decreased-response group (Fig.4f), and baseline FC was inversely related to drug-induced change across individuals (Fig.4g-h). This inverse baseline–ΔFC relationship significantly exceeded a permutation-derived null distribution generated by shuffling drug-induced changes across individuals (10,000 permutations; midazolam: two-sided *P*=0.0056, ketamine: two-sided *P*=0.069, showing a trend in the same direction; see Methods), indicating that the bidirectional response is unlikely to reflect regression to the mean alone. Thus, pharmacological E/I perturbation in vivo reproduces the bidirectional, baseline-dependent modulation of NP circuit predicted in silico by DTBs.

To match the pharmacological dataset (MID only), we restricted virtual modulation to MID-specific NP connectivity to place the model and pharmacological data in the same task context. Importantly, a bidirectional, baseline-dependent response toward virtual modulation of MID-specific NP connectivity was preserved (Supplementary Results; Fig. S13), indicating that the core dynamic property of the NP circuit is robust to this task restriction.

We then asked whether a quantitative mapping learned entirely in silico could generalise to predict individual pharmacological responses. In DTBs, we first estimated a linear mapping between simulated baseline MID FC and virtual modulation-induced changes in NP-related connectivity. Applying this same mapping to empirical placebo MID FC, we obtained subject-specific predictions of ketamine- and midazolam-induced FC changes. Predicted and observed drug-induced changes closely matched at the individual level (Fig.4i-j) and stratified participants with opposite response directions (AUC of ketamine=0.69; AUC of midazolam=0.83; Fig.4k), further supporting the translational relevance of DTB-derived virtual modulation.

Together, these findings show that DTBs not only recapitulate individual-level, baseline-dependent response structure observed under pharmacological perturbation, but also provide a mechanistic template for forecasting individual drug-induced circuit shifts from baseline network configuration.

### DTB-derived perturbational responses predicts future changes in symptoms

We next asked whether DTB-derived perturbational responses carry prospective information about longer-term symptom trajectories. Focusing on internalising symptoms (given the limited prevalence of externalising symptoms at age 23), we first learn a linear mapping from empirical NP connectivity at age 19 to concurrent internalising symptom scores (Fig.4l). We then applied this mapping to simulated NP connectivity before and after virtual E/I modulation to obtain subject-specific predicted symptom changes, termed the “behavioural restoration” index.

We quantified observed symptom change over four years (Δbehaviour, ages 19–23) and tested whether the DTB-derived “behavioural restoration” index predicted these longitudinal changes. Across individuals, baseline symptoms and the restoration index for AMPA modulation together explained ∼23% of the variance in the symptom changes (Fig.4m). The restoration index alone was significantly correlated with empirical symptoms changes (Fig.S14), indicating that individuals whose NP circuit was more “normalisable” in silico were less likely to deteriorate and more likely to remit. Importantly, the restoration index contributed unique predictive value beyond baseline symptoms, significantly increasing explained variance by ∼5% relative to a symptom-only model (Fig.S14). However, the behavioural restoration index for GABA-A modulation was not correlated with empirical symptom changes (Fig. S14).

These findings suggest that DTB-derived perturbational response metrics capture not only cross-sectional heterogeneity but also clinically relevant dynamical properties that forecast longitudinal trajectories. Taken together with the pharmacological validation above, these results position DTB models as a promising tool for predicting and stratifying individualised responses to neurotransmitter-targeted interventions, with potential implications for the development and personalization of future psychiatric treatments.

## Discussion

We simulated a transdiagnostic, behaviourally relevant task-evoked functional network phenotype (i.e., NP factor) within individualised neuronal-scale digital twin brain models. By systematically perturbing AMPA- and GABA-A–mediated synaptic gain, we quantified individual-specific, bidirectional network responses and linked these virtual response profiles to both pharmacological fMRI effects and longitudinal symptom trajectories. These results demonstrate that digital brain models can move beyond descriptive simulation of neural activity to perform controlled in silico perturbations for mechanistic interrogation and individualised counterfactual prediction. By introducing biologically interpretable synaptic parameters, our framework allows precise manipulation of excitation–inhibition balance at the neuronal scale while preserving each individual’s anatomical and task-evoked constraints, establishing a new class of perturbational and predictive digital brain models for computational neuroscience and psychiatry.

The NP factor represents a compact functional connectivity pattern that relates to a broad spectrum of internalising and externalising symptoms across heterogeneous populations. It is characterised by reduced functional integration among limbic, default-mode, frontoparietal control, and cingulo-opercular networks, which jointly support reward processing, cognitive control, and attention allocation^64–67^. Disruptions within and between these networks have been reported across mood, anxiety, and substance use disorders, as well as in population-based samples spanning the full range of symptom severity^18–20,68^. In this sense, the NP factor reflects a shared circuit phenotype rather than a disorder-specific marker. Importantly, we treat NP as a continuous circuit dimension rather than a diagnostic classifier. This makes it suitable as a perturbational readout: it is sufficiently low-dimensional to allow explicit predictions about network reconfiguration, while remaining anchored in identifiable large-scale functional systems.

A central contribution of the NP-DTB framework is that it renders a descriptive task-evoked connectivity phenotype experimentally manipulable. Conventional neuroimaging analyses typically relate connectivity measures to behavioural symptoms, but they do not permit controlled intervention at the circuit level^69,70^. By contrast, DTBs establish an explicit link between large-scale functional network organization and synaptic-scale parameters, enabling in silico perturbation of defined excitatory and inhibitory mechanisms. In addition, DTBs reproduce individualised, task-evoked functional connectivity patterns that are challenging for the conventional whole-brain models, which often rely on coarse-grained regional parcellations and focus primarily on resting-state dynamics^71,72^. This task-state emphasis is critical, as many clinically relevant circuit abnormalities are context dependent^20,73^, and network configurations expressed during reward processing or inhibitory control cannot be inferred from resting-state activity alone^74^. By assimilating task-state signals and applying perturbations at the synaptic level, DTBs generate individual-specific predictions of how functional circuits respond to mechanistically interpretable excitatory or inhibitory modulation.

Using this framework, we observed a pronounced baseline-dependent heterogeneity in NP connectivity response to virtual E/I perturbations. Individuals with lower baseline NP factor scores tend to show larger increases following perturbation, whereas those with higher baseline NP show modest increases or even decreases. Interestingly, both excitatory-leaning (AMPA) and inhibitory-leaning (GABA-A) perturbations often shifted NP connectivity in the same direction within a given individual model. This convergence does not contradict excitation–inhibition balance; rather, it reflects the nonlinear, regime-dependent dynamics of large-scale recurrent networks, in which multiple control parameters can move the system toward a balanced operating regime depending on baseline state^75–77^, consistent with prior empirical and computational modeling studies^57,78–81^. Notably, the bidirectional convergence of excitatory- and inhibitory-leaning perturbations at the network level does not imply equivalence of their molecular mechanisms, but instead reflects compensatory and nonlinear circuit dynamics in which E/I balance acts as a key control variable^57,79,80^.

Two independent lines of evidence further support the biological plausibility of the causal links between microscale neurotransmitter systems and large-scale functional networks inferred by the DTB models. First, PET-derived receptor maps show that regions contributing to the NP circuit are enriched in receptor systems involved in regulating excitation–inhibition balance^46,49,51,52^, providing a neurochemical context for why E/I perturbations plausibly modulate this circuit, although these maps are group-level and cannot establish individual causality. Second, pharmacological fMRI data provide an independent empirical perturbation reference for examining how excitatory and inhibitory neurotransmitter modulation affects NP-related connectivity. Although ketamine and midazolam did not produce robust group-mean effects on NP-related connectivity, individual-level analyses revealed bidirectional, baseline-dependent responses that closely mirror DTB predictions. The convergence between in silico perturbations and pharmacological modulation in vivo suggests that heterogeneity in NP network responses may reflect differences in baseline circuit state, a property also evident in pharmacological responses, rather than an artifact of the digital twin models.

Therefore, a key future implication of the DTB framework is that an individual’s baseline functional network configuration carries information about the direction and magnitude of network change under perturbation. This is supported by our analyses showing a significant inverse relationship between baseline NP connectivity and modulation-induced ΔNP in both AMPA- and GABA-A–mediated virtual perturbations, as well as in pharmacological fMRI data (Fig. 3g–h; Fig. 4g-k). Depending on their initial circuit state, the same intervention may increase network integration in some individuals while decreasing it in others. In this work, we illustrate the translational potential of this approach in two ways. First, we learned a mapping from baseline NP connectivity to virtual perturbation responses in silico and applied it out-of-sample to forecast drug-induced circuit changes in vivo. Second, by linking virtual NP network changes to behaviour, we define a “behavioural restoration” index that quantifies an individual’s brain circuit susceptibility to intervention. This index predicts differences in future symptom trajectories beyond what baseline symptom severity alone can explain, providing a measurable, testable readout of circuit “normalisability” based directly on the model’s simulated perturbation responses. Beyond prediction, the DTB framework also supports mechanistic stratification based on baseline brain network configuration rather than symptom severity alone. Individuals with similar symptom burden but distinct NP configurations can follow divergent trajectories over time, offering a mechanistic explanation for why symptom-based stratification often fails to anticipate treatment response. However, we should note that the excitatory and inhibitory perturbations implemented in the DTB represent controlled shifts in synaptic gain that move the system across distinct network operating regimes, rather than modeling reality drug actions. Thus, this predictive framework is intended for risk stratification and mechanistic insight, rather than deterministic individual-level prognosis.

Notably, model-empirical similarity increased with neuronal resolution in voxel-wise simulations, consistent with scaling properties of the previous studies^9,10^. Regional DTBs showed higher similarity in aggregated BOLD dynamics but lower correspondence in functional connectivity relative to voxel-wise models, likely reflecting reduced spatial granularity for capturing individual-level network structure. Nevertheless, regional models remained computationally tractable and retained sufficient fidelity to predict task performance and empirical group-level differences, supporting their use for population-scale perturbational analyses. Rather than serving as exact replicas of biological brains, DTBs should be viewed as experimentally controllable models that enable causal hypothesis testing in systems otherwise inaccessible to direct manipulation.

Several limitations should be noted. DTB inferences depend on simplifying modelling assumptions, e.g., fixed excitatory-inhibitory neuron ratios, homogeneous synaptic architecture, and simplified neuronal dynamics. Accordingly, AMPA- and GABA-A–labelled parameters should be interpreted as phenomenological gain parameters rather than direct mapping to drug mechanisms. The present DTB framework does not incorporate the full range of neurotransmitter systems, nor cerebellar microcircuit dynamics, and pharmacological validation was limited to a single task and cohort. Although PET receptor maps provide biological plausibility, they do not capture individual neurochemical variability. Despite these constraints, the convergent evidence suggests that a compact task-evoked circuit phenotype can be linked to explicit synaptic control parameters in individualised models, yielding baseline-dependent, and bidirectional predictions that align with pharmacological perturbations and carry prognostic information. Together, these findings motivate a shift from descriptive biomarkers toward perturbational models that explain and predict individual differences in psychiatric trajectories.

## Methods

### Participants and MRI dataset

#### IMAGEN dataset

The discovery dataset used to idenitify brain phenotype of transdiagnostic symptoms was derived from the IMAGEN cohort (https://www.imagen-project.org/), a longitudinal, population-based study comprising behavioural, neuroimaging, environmental, and genetic data from approximately 2,000 participants assessed at ages 14, 19, and 23^1,2^. IMAGEN participants were recruited from eight research sites across the United Kingdom, Germany, France, and Ireland, and were cognitively normal with no history of psychiatric diagnoses. To reduce brain developmental confounds, we focused on early adulthood, specifically the follow-up 2 timepoint (aged 19y) of the IMAGEN study. At this time point, task-fMRI data were available for the Monetary Incentive Delay (MID, n = 1,390) and Stop Signal Task (SST, n = 1,394). After excluding participants with excessive head motion (mean framewise displacement > 0.5 mm) and ensuring matched multimode data, the final sample included 1,050 participants (mean age = 18.41 ± 0.67 years; female/male = 564/486).

The task-fMRI paradigms used in this study are summarized below. The MID task comprised 42 trials across three reward conditions (no-win, small-win, and big-win). Each trial consisted of a reward cue (250 ms), a fixation period (4-4.5 s), a target response, and reward feedback (1,450 ms). The SST included 480 go trials (respond to arrow direction) and 80 stop trials (inhibit response when a stop signal appeared ∼300 ms after the go cue). The Emotional Face Task (EFT) presented angry, neutral, and happy human faces in a passive-viewing paradigm without responses. Each emotional condition comprised four trials, with each trial lasting 18 s. More details were available in the original paper of IMAGEN study ^2^.

#### STRATIFY dataset

We validated brain phenotype of transdiagnostic symptoms in independent samples from the STRATIFY cohorts^3^, which used identical MRI protocols and behavioural assessments to IMAGEN. These datasets included patients with psychiatric diagnoses and demographically matched healthy controls. After the same screening procedure with IMAGEN, the final sample included 209 patients (age = 22.26 ± 2.21; female/male = 135/74) and 225 healthy controls (age = 21.40 ± 1.72; female/male = 129/96). Specific diagnoses included alcohol use disorder (AUD, female/male = 58/40), major depressive disorder (MDD, female/male = 75/29), psychosis (female/male = 2/4), and attention-deficit/hyperactivity disorder (ADHD, female/male = 0/1) for STRATIFY.

Clinical diagnoses were determined based on established thresholds from standardized assessment tools consistent with DSM-5 criteria. Individuals with AUD were defined by an Alcohol Use Disorders Identification Test (AUDIT)^4^ score greater than 15. MDD was defined by a Patient Health Questionnaire-9 (PHQ-9)^5^ score greater than 15. Psychosis was defined using ICD-10 diagnosis, with inclusion requiring at least one episode of schizophreniform or chronic schizophrenia. Healthy controls were screened using rigorous inclusion criteria to minimize confounding factors. Participants were excluded if they had: a) any current or past mental health disorder^6^, b) first- or second-degree relatives with mental health diagnoses, c) regular use of medication for serious physical health conditions, d) learning difficulties (e.g., Wechsler Adult Intelligence Scale verbal comprehension score below the 10th percentile), or e) any history of recreational drug use.

#### Pharmacological dataset

To validate the effects of in sillco neurotransmitter manipulation in vivo, we utilized pharmacological task-fMRI data^7^ from 30 healthy male participants (mean age 27.3 ± 6.2 years). Each participant underwent three separate scanning sessions in a randomized, single-blind, placebo-controlled, three-way crossover design, receiving ketamine, midazolam, or placebo. A minimum 48-hour interval was maintained between sessions to prevent carryover effects. The experimental procedure included a 16-minute resting-state fMRI scan, during which drug administration began at the 7-minute mark, followed by task-fMRI acquisition during the MID task. Three participants were excluded due to missing MID behavioural recording files (n = 2) and T1-weighted structural MRI data (n = 1). The MID task used here comprised 90 trials across two conditions: no-win (45 trials) and win (45 trials). Details on administration protocols are available in the original study^7^.

All above studies received approval from the local ethics committee, and written informed consent was obtained from all participants.

### Behavioural symptoms assessments

We assessed behavioural symptoms for each individual from IMAGEN and STRATIFY cohorts using the Development and Well-Being Assessment^8^ (DAWBA) and Strengths and Difficulties Questionnaire^9^ (SDQ). To characterise externalising and internalising symptoms, we summed the scores of relevant items from DAWBA and SDQ^3^. The externalising symptoms included ADHD (8 items) and conduct disorder (CD, 7 items), while the internalising symptoms encompassed eating disorder (ED, 5 items), specific phobia (SP, 13 items), general anxiety disorder (GAD, 7 items), and depression (DEP, 8 items). A detailed list of the specific items is provided in Table S2.

### MRI acquisition and preprocessing

The structural and functional MRI protocols for the IMAGEN and STRATIFY studies were harmonised across sites and scanner manufacturers (Siemens: 6 sites, Philips: 2 sites, General Electric: 1 site, and Bruker: 1 site), Standardized hardware for visual and auditory stimulus presentation (Nordic Neurolabs, Norway) was used at all sites.

#### Structural T1-weighted images

T1-weighted images (T1-w) of the IMAGEN and STRATIFY cohorts were acquired using a protocol based on those from the ADNI study (https://adni.loni.usc.edu/data-samples/adni-data/neuroimaging/mri/mri-scanner-protocols/). Several parameters in these protocols deliberately differ between scanner models, in order to produce consistent image contrast and quality despite implementation differences between manufacturers; all scans were however collected with a sagittal slice plane and voxel size = 1.1 × 1.1 ×1.1 mm^3^.

The structural T1-w images were preprocessed using fMRIPrep^10^ (version 20.2.3), a robust and standardized pipeline based on Nipype, supported by Advanced Normalization Tools (ANTs, version 2.3.3), FMRIB Software Library (FSL, version 5.0.9), Analysis of Functional NeuroImages (AFNI, version 20160207), and FreeSurfer (6.0.1) packages. Anatomical preprocessing included bias field correction, skull-stripping, tissue segmentation and nonlinear registration to standard Montreal Neurological Institute (MNI) space.

#### Diffusion-weighted images

Diffusion-weighted images (DWIs) of IMAGEN and STRATIFY cohorts were acquired using a 2D diffusion weighted EPI sequence: TR = 15,000 ms; TE = 104 ms; FOV = 307 × 307 mm^2^; number of slices = 60; scan time = 9 min 45 s; voxel size = 2.4 × 2.4 × 2.4 mm^3^; 36 optimal non-colinear diffusion-weighted directions with b = 1,300 s/mm^2^.

The structural DWIs were preprocessed using the FSL and MRtrix3 package. Eddy current-induced distortions and head motion were corrected using FSL’s eddy tool, and the rotated b-vectors were updated using eddy_rotated_bvecs. The preprocessed DWI data was then converted to the MRtrix3 format (.mif) with the mrconvert tool. The mean b0 image was extracted using the mrmath command (mean), followed by brain extraction with the bet tool (fractional intensity threshold = 0.3) to generate a brain mask. The white matter fiber orientation distributions (FODs) within each voxel were estimated using constrained spherical deconvolution (CSD), with response function estimation performed using the Tournier method. The anatomical T1-weighted image was registered to the native diffusion space using FSL’s flirt tool. A nonlinear transformation to MNI space was then computed using FSL’s fnirt, and the inverse warp was generated with invwarp. The resulting warp was subsequently converted into an MRtrix-compatible format for fiber transformation. A five-tissue-type segmentation was carried out with the 5ttgen command. The gray matter-white matter interface (GMWMI) was extracted using the 5tt2gmwmi tool to serve as the seeding region for tractography. Whole-brain fiber tracking was performed using the tckgen command with the iFOD2 algorithm, seeding from the extracted GMWMI and applying Anatomically Constrained Tractography (ACT). The generated fiber tracks were transformed from diffusion space to MNI space and generate voxel-wise and regional-level structural connectivity matrices.

#### Resting-state and task-state functional MRI images

The resting-state fMRI scans of IMAGEN and STRATIFY cohorts were acquired using a 2D gradient echo EPI sequence with the following parameters: TR = 2,200 ms; TE = 30 ms; FA = 75°; FOV = 220 × 220 mm^2^; number of slices = 40; number of volumes = 164; and voxel size = 3.4 × 3.4 × 2.4 mm^3^. Task-state fMRI scans were conducted with identical scanning parameters, except for the number of volumes, that is 191 volumes for the MID task, 349 volumes for SST task, and 202 volumes for EFT task.

Addtionally, the pharamacological MID task-state fMRI were acquired on a 3T scanner (Siemens Skyra, Erlangen, Germany), using the EPI sequence: TR = 2,200 ms; TE = 27 ms; FA = 79°; FOV = 215 × 215 mm^2^; number of slices = 30; number of volumes = 410; and voxel size = 3 × 3 × 3 mm^3^.

All fMRI data were preprocessed using fMRIPrep (version 20.2.3), including motion correction, slice timing correction, and susceptibility distortion correction using fMRIPrep’s fieldmap-less approach. Functional MRI images were co-registered to the T1-weighted image using boundary-based registration and then normalized to MNI152 space. Confound regressors were extracted to control for physiological and motion-related artifacts, including framewise displacement, DVARS, and CompCor components. All transformations were applied in a single interpolation step to minimize resampling effects. Then the preprocessed data underwent temporal detrending to remove low-frequency drifts, followed by spatial smoothing using a full-width at half maximum (FWHM) of 6 mm Gaussian kernel and a band-pass filtering (0.01 ∼ 0.1 Hz). Quality control was performed by excluding subjects with incomplete demographic or scanning timepoints; excessive head motion (mean framewise displacement > 0.5 mm); or unsuccessful spatial normalization.

### Identification of a compact network phenotype of psychopathology in IMAGEN

Followed by previous work identifing the neuropsychopathology (NP) factor in the IMAGEN baseline at age 14^3^, we constructed MID and SST task-specific functional connectivity (FC) matrics using the CONN toolbox (version 16.h) and a well-established 268-node whole-brain functional parcellation^11^, applying weighted generalised linear models after regressing out task condition regressors and nuisance variables. Subsequently, we employed a Connectome-based Predictive Modeling (CPM) ^12^ with a 50-fold cross-validation to predict each behavioural symptom scores (including ADHD conduct disorder, eating disorder, specific phobia, general anxiety disorder, and depression) from each task-specific functional connectomes (SST conditions included stop-success, stop-failure, and go-wrong; MID conditions included positive feedback, negative feedback, and reward anticipation). The model performance was assessed using the Spearman’s correlation between predicted and empirical behavioural symptoms scores. This procedure was repeated 1,000 times, and only edges selected in over 95% of models were used for further analysis. The mean *P* values across 1,000 repetitions were corrected for multiple comparisons across 36 predictive models using false discovery rate (FDR) correction (*q* < 0.05). Task-specific functional connectivity (FC) matrices were then selected if they showed statistically significant prediction performance for at least three behavioural symptom domains, ensuring robustness across multiple diagnostic categories.

From these selected task-specific FC matrices, we extracted edges that significantly predicted both externalising and internalising symptoms. Edges that were consistently associated with behavioural symptoms across all selected task conditions were defined as transdiagnostic associated edges. To improve interpretability, we stratified these transdiagnostic associated edges into positive or negative FC profiles based on the direction of their associations with behavioural symptoms. The summed strengths of these two profiles were termed as positive and negative NP scores, respectively.

### Case-control comparison of NP factors in STRATIFY

To test whether the NP scores generalise to clinical populations, we re-extracted these transdiagnostic associated FC profiles in the independent STRATIFY dataset and re-calculated NP scores for each individual. We then compared NP scores between patients and healthy controls, as well as among depression and alcohol use disorder subgroups, using independent-samples *t* tests while covarying for sex, site, and head motion (mean framewise displacement). Multiple comparisons were corrected using the Bonferroni method.

### Permutation testing of PET receptor maps enrichment in NP-related regions

To assess the neurochemical context of the NP factor, we examined whether the NP-related regions exhibited higher receptor map values relative to non-NP regions using published whole-brain PET-derived receptor density maps^13^ (https://github.com/netneurolab/hansen_receptors/tree/main/data/PET_nifti_images). For a given PET map, we first extracted regional expressive levels using the 268-node functional parcellation^11^, and then computed the empirical difference in mean values between NP (n = 32) and non-NP (n = 236) regions. Statistical significance was assessed using permutation testing (10,000 times), in which region labels (NP-related vs NP-unrelated) were randomly reassigned while preserving the original group sizes. For each permutation, the difference in mean receptor values between the permuted groups was recomputed, yielding a null distribution expected under no spatial specificity. Two-sided permutation *P* values were calculated as the proportion of permuted differences whose absolute value exceeded the empirical difference. Resulting *P* values were corrected for multiple comparisons across 39 receptor maps using FDR correction.

### Multimode neuroimaging data for Digital Twin Brain (DTB) models

To constructing individualised DTB models, we extracted voxel-wise multimodel neuroimaging data, including grey matter volume, white matter structural connectivity, and functional BOLD signals from resting-state and task-state fMRI.

Gray matter volume was estimated using voxel-based morphometry (VBM) in SPM12 (Matlab R2020b). T1-weighted images were segmented into tissue maps, normalized to MNI space using DARTEL, modulated by Jacobian determinants, and smoothed with an 8 mm FWHM Gaussian kernel at 3 × 3 × 3 mm³ resolution.

White matter structural connectivity was derived from whole-brain tractography using the iFOD2 algorithm in MRtrix3, with anatomical constraints (ACT). Tracking used 5 million streamlines seeded from the GMWMI, with a step size of 0.2 mm, curvature threshold of 45°/step, and length range of 3–250 mm.

To extract BOLD signals, we first generated individualised brain mask by selecting voxels that: a) overlapped with both the 268-node functional parcellation^11^ and the MNI152 template, b) had existing white matter structural connections, and c) were located in the cortex or subcortex. On average, ∼12,000 voxels per subject were retained. The time series from these voxels were then extracted as simulation targets for the DTB models.

Importantly, incorporating specific neuronal models for the cerebellum and brainstem would substantially increase model complexity and computational cost, particularly given that the DTB simulations are implemented at the scale of hundreds of millions to billions of neurons. We therefore adopted a parsimonious modeling strategy focused on cortical and subcortical circuits, while acknowledging that this exclusion represents a limitation and that future extensions incorporating differentiated regional parameterization may enable more comprehensive characterization of NP network dynamics.

### Construction of individualised task-state DTB models

Followed by the framework developed by Lu et al.^14,15^, we firstly constructed a excitation-inhibition balanced spiking neural network constrained by individual multimode neuroimaging data. In this network, each voxel was modeled as a sub-unit comprising one excitatory and one inhibitory neuronal population. Each voxel included both excitatory and inhibitory neurons at a fixed ratio of 4:1^16^. The number of neurons within each voxel was determined by its proportional grey matter volume. Within-voxel synaptic architecture followed a 4:1:2 ratio of internal excitatory, internal inhibitory, and external excitatory synapses, based on anatomical organization of the cat visual cortex^17^. Between-voxel connections were weighted by row-normalized structural connectivity derived from diffusion MRI, and exclusively excitatory^18,19^. Neuronal activity was simulated using the leaky integrate-and-fire model^20^, incorporating α-amino-3-hydroxy-5-methyl-4-isoxazolepropionic-acid (AMPA; excitatory-leaning) and γ-aminobutyric-acid-A (GABA-A; inhibitory-leaning) synaptic conductances to represent glutamatergic and GABAergic signaling:

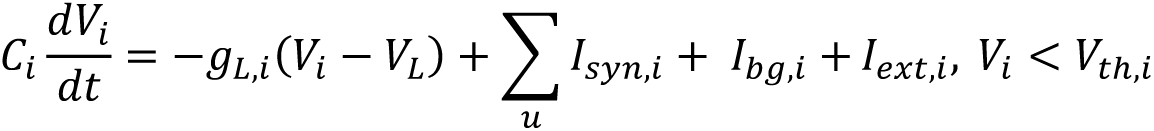

where *C_i_* is the capacitance of the neuron membrane, *g_L,i_* is the leakage conductance, *V_i_* is the membrane potential of neuron i, *V_L_* is the leakage voltage, *I_syn,i_* is the synaptic currents of two synapse types (AMPA and GABA-A), *I_bg,i_* is the background current as noises, and *I_ext,i_* is the external current input for the task. It should be noted that the *I_ext,i_* are the independent parameters, which are estimated using empirical task-fMRI data. The background noise is given by independent Ornstein–Uhlenbeck processes and described as in the original paper^15^.

Simulated neuronal activity was quantified as mean firing rate of neuronal population within each voxel, and was transformed into simulated BOLD signals using the Balloon-Windkessel model^21^. The model performance was quantified by Pearson’s correlation between simulated and empirical task-state BOLD signals.

To optimize the fit between simulated and empirical BOLD signals, we applied the Hierarchical Mesoscale Data Assimilation (HMDA) method^15^ to estimate voxel-wise, time-dependent hyperparameters. This assimilation procedure iteratively alternated between forward simulating neuronal activity and parameter updating to reduce the mismatch between simulated and observed BOLD signals. At each time point for each assimilated voxel, the parameters of synaptic conductances for AMPA and GABA-A were sampled from gamma distributions, which were constrained by voxel-specific hyperparameters. These synaptic conductances governed the simulated neuronal dynamics, which were transformed into BOLD signals. Hyperparameters were then iteratively updated so that the resulting simulated BOLD signals progressively converged toward the empirical fMRI data. Parameter updating was implemented using a diffusion ensemble Kalman filter (EnKF), which estimates latent states and parameters by propagating an ensemble of 30 parallel model realizations. More methodological details of HMDA are describedin Lu et al.’s original paper^15^.

Resting-state digital twin brain models were first obtained by assimilating empirical resting-state fMRI BOLD signals for all voxels using the HMDA framework. This procedure yielded individualised resting-state DTBs with converged hyperparameters and stable neuronal dynamics. To construct task-state DTB models, we next assimilated empirical task-state fMRI BOLD signals from task-engaged regions to infer digital external input currents. These inferred digital input currents were then injected into the corresponding voxels of the resting-state DTB to induce task-related neuronal activity. For regions not directly engaged by the task, model hyperparameters were fixed to the mean values estimated during the resting-state assimilation, ensuring a stable baseline while allowing task-specific dynamics to propagate through the network.

Task-relevant regions were identified based on prior literature and meta-analytic activation maps from the Neurosynth database. For the MID task, regions where received assimilated currents included the orbitofrontal cortex, dorsal anterior cingulate cortex, insula, amygdala, dorsal striatum, and nucleus accumbens, associated with reward processing. For the SST task, assimilated regions involved the orbitofrontal cortex, anterior and dorsolateral prefrontal cortex, dorsal ACC, insula, and thalamus, related to executive control. For the EFT task, assimilated regions included the amygdala, dorsal striatum, hippocampus, insula, thalamus, anterior and posterior cingulate cortex, dorsolateral prefrontal cortex, and orbitofrontal cortex, which are related to emotional recognition and processing. Primary visual were included for all three tasks, and sensorimotor cortices for first two tasks. A full list of assimilated regions is provided in Table S6.

### Assessment of subject-specific information in DTB-assimilated hyperparameters

To evaluate whether DTB-assimilated hyperparameters capture subject-specific neural dynamics independent of empirical data, we analyzed EFT task-based fMRI data from an independent subset of four male participants, matched for scanning site (Berlin) and age (one MDD and one AUD, both aged 21 years, and two HCs aged 22 years). Each participant completed four trials per emotional block (angry, neutral, and happy), yielding 12 trials per subject.

We first independently estimated hyperparameters for each trial, without using any other trial of the same participant. Each trial-specific hyperparameter was then injected into noise-initialized DTB models to generate simulated BOLD time series of matched duration. Trial-level activation maps were computed from simulated and empirical BOLD signals using identical preprocessing and statistical modeling procedures. To quantify subject specificity, we correlated activation maps from all simulated and empirical trials across subjects using Pearson correlation coefficients, yielding a 48 × 48 similarity matrix (4 subjects × 12 trials), and compared correlations between simulated and empirical trials from the same individual (self–self) with those between trials from different individuals (self–other) using a Mann–Whitney *U* test.

In addition, we tested whether simulated activation maps, generated without re-assimilation of empirical data, were sufficiently distinctive to support individual identification. For each participant, we created subject-specific empirical templates by averaging the first two trials of each emotional condition. Hyperparameters estimated from these first-half trials were used to simulate BOLD time series for the remaining trials without assimilation. We then correlated activation maps from simulated second-half trials (4 subjects × 6 trials) with all subject-specific templates, and aligned each simulated trial to the subject with the highest correlation. The same identification procedure was applied to empirical activation maps from the second-half trials as a benchmark.

### Calibration experiment of DTB model scales

The DTB framework allows flexible specification of total neuronal count across a wide range, from approximately 1 million to 86 billion neurons^14,15^. To balance biological fidelity, perturbational controllability, and computational feasibility, we performed calibration experiments to determine optimal model resolutions for three complementary analytical purposes: high-fidelity simulation, controlled perturbational exploration, and population-level inference.

We first evaluated the influence of model scale using the DTB of an illustrative healthy control participant (HC01), by varying neuronal counts from 10 million to 5 billion and assessing simulation fidelity across resolutions. Model performance was quantified using the mean voxel-wise Pearson’s correlation between simulated and empirical BOLD signals, as well as correlations between simulated and empirical whole-brain functional connectivity matrices (268-node functional parcellation^11^). These experiments established the relationship between neuronal scale and task-state simulation accuracy, providing an empirical basis for selecting biologically realistic yet computationally tractable model resolutions.

Second, to identify the optimal resolution for the population-scale simulations, we constructed reduced-scale DTBs ranging from 1 to 9 million neurons for the same participant. In these models, individual MRI data were represented at the regional rather than voxel-wise level. The number of neurons per region was assigned in proportion to the regional grey matter volume. While voxel-wise grey matter volumes could in principle be used, the total number of neurons in millions of neuron models would make it impractical to allocate neurons to every individual voxel, as some voxels have a minimal ratio of grey matter volume. By using regional-level allocation, we ensured that the neuron distribution faithfully reflected grey matter proportions while remaining computationally tractable. Between-regional connections were weighted by row-normalized regional structural connectivity derived from diffusion MRI. The other details were consistent with the methodology used in our voxel-wise simulation models. Model performance was quantified using the mean regional-level Pearson’s correlation between simulated and empirical BOLD signals, as well as correlations between simulated and empirical whole-brain functional connectivity matrices (268-node functional parcellation^11^).

Together, these calibration procedures ensured that DTB simulations maintained high fidelity while enabling scalable individualised and population-level analyses.

### DTB Simulations for four illustrative participants

Following the calibration experiments described above (see Supplementary Results), high-fidelity 1-billion-neuron DTBs were constructed for four illustrative participants: two patients (one MDD and one AUD) and two healthy controls (HC01 and HC02; demographics information in Table S7). The patients were selected as those with the highest negative NP scores within their respective diagnostic groups, indicating the highest symptom burden. Among the healthy controls, HC01 and HC02 were chosen from the subset of participants with the lowest behavioural symptom summed scores in the IMAGEN cohort. HC01 had the lowest negative NP score, whereas HC02 had the highest negative NP score, illustrating the range of NP values even among individuals with minimal behavioural symptoms. All selected participants had complete multi-modal imaging data, including T1-weighted, DTI, and fMRI scans.

In addition to billion-neuron simulations, 100-million-neuron DTBs were constructed for the same participants to enable computationally efficient perturbational analyses while retaining individualised model characteristics. This intermediate resolution was selected by calibration results showing no clear performance inflection point across neuronal scales (Fig.S4; Table S8), suggesting that reduced-scale models could preserve essential network properties while improving controllability for systematic parameter exploration.

To evaluate whether 100-million-neuron DTBs retained subject-specific network phenotypes, we extracted simulated NP factor scores from each individualised model and compared them with empirical NP factors derived from all individuals in the IMAGEN and STRATIFY datasets. Similarity was quantified using mean-squared error (MSE) between each pair of simulated and empirical NP factors. For each illustrative participant, self-to-self MSE (simulated versus own empirical NP factor) was contrasted against cross-participant MSE (simulated versus all other empirical scores). Statistical significance was evaluated using permutation testing across all 1,575 simulated - empirical pairings.

### Systematic parameter sweeps of virtual AMPA and GABA-A perturbations

Based on 100-million-neuron DTB models of four illustrative participants, we performed systematic perturbations of AMPA- and GABA-A-mediated synaptic conductances. AMPA conductance was varied from 0.0020 to 0.0052 in steps of 0.0004 (baseline = 0.0008), and GABA-A conductance from 0.0015 to 0.0040 in steps of 0.0005 (baseline = 0.0015). The baseline levels of AMPA and GABA-A in the model were determined across two key dimensions: the mean firing rate of the neuronal population (<10 Hz), consistent with physiological observations^22^, and the synchrony measure, which represents the degree of oscillatory behaviour. These metrics identified the optimal parameter window where the model maintains robust, reasonable neuronal firing while avoiding excessive synchronization^23^, thereby ensuring a stable and biologically plausible baseline for the model.

Crucially, the upper limit for the parameter of AMPA conductance was constrained by the stability of the simulated BOLD signal: beyond a specific threshold, excessive neuronal firing led to a cessation of dynamic fluctuations in the BOLD signal. We considered such states as “over-firing” regimes, where the hemodynamic response reaches a non-biological plateau and no longer reflect meaningful neural dynamics^24^. During GABA-A manipulations, AMPA conductance was held fixed at the value that had produced the maximal NP factor enhancement during prior AMPA sweeps, allowing controlled assessment of excitatory-inhibitory balance, and preventing excessive inhibition that could silence neuronal activity and abolish intrinsic BOLD fluctuations^25^.

These synaptic conductance parameters were implemented as global control variables, applied uniformly across the whole-brain model rather than specific regions. Importantly, they were directly applied to DTB models that had already been calibrated to individual empirical fMRI data; therefore, no additional data assimilation or parameter refitting was required during virtual perturbations. The NP network responses were obtained by forward simulation under systematically varied synaptic levels. This design allows direct assessment of perturbational sensitivity around individualised baseline operating points. To ensure reproducibility, each perturbation was repeated five times, and the mean NP factor scores across repetitions were used for subsequent analyses. Finally, while multiple combinations of synaptic parameters can give rise to similar network outputs, our goal was not to recover exact physiological parameter values, but to use biologically interpretable synaptic conductance parameters as control variables to probe regime-dependent network responses.

### Evaluation of cross-task generalisability of virtual perturbations

To test whether the effects of AMPA- and GABA-A–mediated perturbations on the NP factor generalise across cognitive contexts, we conducted the same virtual perturbation to the DTB models that simulate EFT task activity of the same four illustrative participants. Specifically, we first calibrated DTB models at a 100-million-neuron scale to simulate BOLD signals of EFT task-based fMRI data, and then applied the optimal perturbational parameters identified in the parameter sweeps to these DTB models. For each participant, NP-related functional connectivity was extracted from baseline and perturbed simulated EFT BOLD signals using the same computational pipeline as the original MID and SST tasks, enabling direct comparison of perturbation effects across cognitive tasks.

### Population-scale virtual perturbations

To assess whether virtual E/I modulations in DTB models generalise across the cohort, we first constructed DTB models for 290 participants, including 72 patients with MDD (age = 22.33 ± 2.20, female/male = 48/24), 59 with AUD (age = 22.41 ± 2.02, female/male = 37/22), and 69 healthy controls (age = 21.45 ± 1.38, female/male = 38/31) from STRATIFY dataset, as well as 90 subclinical participants with high behavioural symptom scores (sum of six externalising and internalising symptoms ≥ 20) from IMAGEN (age = 18.42 ± 0.65, female/male = 74/16). We then applied the optimal perturbational parameters identified in the parameter sweeps to these DTB models.

For each participant, we calculated simulated NP factor scores derived from baseline and perturbed BOLD signals. Group differences in empirical, simulated, and perturbed NP factors were assessed using ANOVA analysis and post-doc tests, controlling for sex, site, and head motion (mean framewise displacement). The paired-sample *t* tests were used to compare simulated and empirical, and simulated and perturbed NP factors, among subgroups. Multiple comparisons were corrected using the Holm-Bonferroni method.

### Prediction of task performance from DTB simulations

To assess the behavioural validity of digital twin brain simulations, we evaluated whether simulated task-state functional connectivity metrics, derived from population-scale simulations, significantly predicted individual differences in task performance during the MID task.

Analyses were conducted in the 287 participants with 3-million-neuron DTB simulations and MID task behavioural data. Individualised task-state functional connectomes were extracted separately for the anticipation-hit and feedback-hit stages of the MID task. Behavioural performance was quantified as mean response time (RT) under three incentive conditions: big-win, small-win, and no-win. Prediction of behavioural performance was performed using a connectome-based predictive modeling framework with 10-fold cross-validation. For each fold, linear models were trained on task-state FC matrices from nine folds to predict RT measures and evaluated on the held-out fold. Prediction accuracy was quantified as the Spearman’s correlation between predicted and observed RT values. The entire cross-validation procedure was repeated 100 times to ensure stability, and prediction accuracies were averaged across repetitions. Separate models were constructed for each combination of task stage (anticipation-hit, feedback-hit) and behavioural measure (big-win, small-win, no-win), resulting in six independent prediction models.

We also examined the prediction performances of empirical task-state FC for establishing a benchmark. Prediction accuracies obtained using empirical FC and DTB-simulated FC were compared using paired-sample *t* tests across the six model combinations.

### Stratification of NP network responses to virtual perturbations

In the population-level analysis, we quantified individual responses to virtual neurotransmitter perturbations in the DTB models. For each participant, we computed the change in simulated NP factor following AMPA and GABA-A modulation relative to baseline. Participants were classified as “increasers” (ΔNP>0 following both modulations) or “decreasers” (ΔNP ≤ 0 for any given perturbation). We compared the distribution of response types across diagnostic groups (patients, high-symptom, and healthy controls), sex, and site using Chi-square tests.

We also assessed the relationship between baseline simulated NP factor and the AMPA- or GABA-A–induced changes using Pearson’s correlation. Statistical significance was assessed using permutation testing (10,000 permutations), in which ΔNP values were randomly reassigned across individuals to generate a null distribution. For each permutation, the correlation between simulated baseline NP and permuted ΔNP was recomputed. Two-sided permutation *P* values were calculated as the proportion of null correlations whose absolute value exceeded the observed correlation, directly testing whether modulation-induced changes reflect systematic baseline dependence rather than symmetric fluctuations or regression-to-the-mean effects.

Finally, we compared simulated baseline NP factors and behavioural symptom measures using Mann–Whitney *U* tests, given unequal subgroup sizes and non-normal distributions. All analyses controlled for sex, scanning site, and mean framewise displacement. Behavioural measures were derived from the 69 entry items of the DAWBA and the SDQ.

### Stratification of individual pharmacological responses

To validate DTB-predicted effects of virtual AMPA and GABA-A modulations on the NP factor, we analyzed an independent pharmacological fMRI dataset in which healthy participants completed MID task-fMRI under placebo, ketamine, and midazolam in a randomised cross-over design. For each participant and drug condition, we computed NP-related MID FC strengths (six edges) and summed them as a “summed MID FC” metric.

We first tested group-level drug effects by comparing the summed MID FC between placebo and drug conditions using paired-sample *t* tests. To further capture individual variability, we computed the drug-induced change in summed MID FC relative to placebo for each participant and applied k-means clustering (500 iterations) to identify two response subgroups. Within each subgroup, paired-sample *t* tests, controlling for head motion, were used to compare placebo and drug conditions. Wilcoxon signed-rank tests were applied when normality or variance assumptions were violated.

We also compared summed MID FC under placebo between response subgroups using a independent-samples *t* test. Finally, the relationship between baseline summed MID FC in placebo and drug-induced FC change across all participants was assessed using Pearson’s correlations. Statistical significance was assessed using permutation testing (10,000 permutations), in which Δsummed MID FC values were randomly reassigned across individuals to generate a null distribution. For each permutation, the correlation between baseline summed MID FC and permuted Δsummed MID FC was recomputed. Two-sided permutation *P* values were calculated as the proportion of null correlations whose absolute value exceeded the observed correlation, directly testing whether drug-induced changes reflect baseline dependence rather than regression-to-the-mean effects.

### Prediction of pharmacological responses from DTB-derived virtual perturbations

To test whether DTBs capture systematic relationships between baseline network configuration and drug-induced NP-related connectivity changes, we first constructed a linear regression model linking simulated baseline summed MID FC and FC changes following virtual AMPA and GABA-A modulations. We then applied this regression model to predict empirical FC changes under ketamine or midazolam from each participant’s baseline FC under placebo. Here, we chose a linear mapping to minimize overfitting and maximize interpretability given the modest sample size. Prediction accuracy was quantified by Pearson’s correlation between predicted and observed FC changes, with significance assessed via permutation tests (1,000 permutations of subject labels).

To evaluate whether DTB-based predictions can distinguish individuals with opposite drug responses, participants were classified as “increasers” or “decreasers” based on the their observed drug-induced change in summed MID FC. The predicted probabilities from the DTB-derived regression model were then used to calculate the area under the receiver operating characteristic curve (AUC) as a measure of how well the model correctly identifies individuals in each response category.

### Prediction of longitudinal symptom changes from DTB-derived virtual perturbations

To test whether DTB-derived virtual E/I modulations predict longitudinal symptom trajectories, we first estimated an empirical linear mapping between NP factors and behavioural symptoms using baseline data. Specifically, we fitted a linear regression linking NP factors at age 19 to summed internalising symptom scores at age 19 across four domains (eating disorder, depression, generalized anxiety, and specific phobia). Externalising symptoms (ADHD and conduct disorder) were excluded due to their low prevalence at follow-up (age 23).

Baseline NP factors and NP factors under virtual AMPA and GABA-A modulation were simulated for the full cohort using 3-million-neuron DTBs. From this cohort, we selected individuals with imaging data at age 19 and symptom assessments at both ages 19 and 23. The empirical NP–symptom mapping derived at baseline was then applied to the DTB-simulated NP factors to generate predicted symptom scores at baseline and after virtual modulation.

For each individual, we defined a DTB-derived “behavioural restoration” index as the difference between predicted baseline and post-modulation symptom scores. Longitudinal symptom change over four years (age 23 minus age 19) was subsequently modeled using multiple linear regression, with baseline symptom scores and the DTB-derived “behavioural restoration” index included as predictors. Prediction accuracy was quantified by Pearson’s correlation between predicted and observed behavioural symptom changes. Model fit was quantified using the coefficient of determination (*r²*). The incremental variance explained by the DTB-derived index (Δ*r²*) was assessed by comparing full and reduced models using *F* tests. Statistical significance of DTB-related effects was further evaluated using 5,000 permutation tests for both *r²* and the regression coefficient associated with the DTB index.

## Acknowledgments

This work received support from the following sources: National Natural Science Foundation of China (32400935 to YX, W2541022 to GS, 82150710554 to GS), National Science and Technology Major Project (2025ZD0215100 to GS), 111 Project (B18015 to JF), Shanghai Municipal Science and Technology Major Project (2018SHZDZX01 to JF), the European Union-funded FP6 Integrated Project IMAGEN (Reinforcement-related behaviour in normal brain function and psychopathology) (LSHM-CT-2007-037286 to GS), the Horizon 2020 funded ERC Advanced Grant ‘STRATIFY’ (Brain network based stratification of reinforcement-related disorders) (695313 to GS), Horizon Europe ‘environMENTAL’, grant no: 101057429 to GS, UK Research and Innovation (UKRI) Horizon Europe funding guarantee (10041392 and 10038599) to SD, Human Brain Project (HBP SGA 2, 785907, and HBP SGA 3, 945539) to GS. The German Center for Mental Health (DZPG), the Bundesministerium für Bildung und Forschung (BMBF grants 01GS08152; 01EV0711; Forschungsnetz AERIAL 01EE1406A, 01EE1406B; Forschungsnetz IMAC-Mind 01GL1745B), the Deutsche Forschungsgemeinschaft (DFG project numbers 458317126 [COPE] to GS, 186318919 [FOR 1617], 178833530 [SFB 940], 386691645 [NE 1383/14-1], 402170461 [TRR 265], 454245598 [IRTG 2773]), the Medical Research Foundation and Medical Research Council (grants MR/R00465X/1 and MR/S020306/1) to SD, the National Institutes of Health (NIH) funded ENIGMA-grants 5U54EB020403-05, 1R56AG058854-01 and U54 EB020403 as well as NIH R01DA049238, the National Institutes of Health, Science Foundation Ireland (16/ERCD/3797) to ALWB and RW.

## Consortia

### DTB Consortium

Yubin Bao, Boyu Chen, Siming Chen, Zhongyu Chen, Fei Dai, Weiyang Ding, Xin Du, Jianfeng Feng, Yubo Hou, Mingda Ji, Peng Ji, Chong Li, Chunhe Li, Xiaoyi Li, Yuhao Liu, Wenlian Lu, Zhihui Lv, Hengyuan Ma, Yang Qi, Edmund Rolls, He Wang, Huarui Wang, Jiexiang Wang, Shouyan Wang, Ziyi Wang, Yunman Xia, Shitong Xiang, Chao Xie, Xiangyang Xue, Leijun Ye, Longbin Zeng, Tianping Zeng, Chenfei Zhang, Jie Zhang, Nan Zhang, Wenyong Zhang, Yicong Zhao & Qibao Zheng

### IMAGEN Consortium

Tobias Banaschewski, Gareth J. Barker, Arun L.W. Bokde, Rüdiger Brühl, Sylvane Desrivières, Herta Flor, Penny Gowland, Antoine Grigis, Andreas Heinz, Herve Lemaitre, Jean-Luc Martinot, Marie-Laure Paillère Martinot, Eric Artiges, Frauke Nees, Dimitri Papadopoulos Orfanos, Tomáš Paus, Luise Poustka, Michael N. Smolka, Sarah Hohmann, Nilakshi Vaidya, Henrik Walter, Robert Whelan, Paul Wirsching & Gunter Schumann

### STRATIFY Consortium

Nilakshi Vaidya, Zuo Zhang, Lauren Robinson, Jeanne Winterer, Yuning Zhang, Gareth J Barker, Arun L.W. Bokde, Rüdiger Brühl, Hedi Kebir, Hervé Lemaître, Frauke Nees, Dimitri Papadopoulos Orfanos, Ulrike Schmidt, Julia Sinclair, Robert Whelan, Henrik Walter, Sylvane Desrivières & Gunter Schumann

### EnvironMENTAL Consortium

Tobias Banaschewski, Sylvane Desrivières, Andreas Heinz, Frauke Nees, Nilakshi Vaidya, Henrik Walter, Vince D. Calhoun & Gunter Schumann

## Author contributions

Conceptualization: G.S., J.F., V.J., W.L., Y.X., S.P.. Methodology: W.L., Y.X., S.P., C.X., S.X., S.Pet., Y.L., J.W., L.Z., H.W., Q.Z., and DTB Consortium. Investigation: Y.X., S.P., S.X., Z.L.. Data acquisition: J.D., J.H., S.M., A.F., T.B., G.J.B., A.L.W.B., R.B., S.D., H.F., P.G., A.G., A.H., H.L., J.-L.M., M.-L.P.M., E.A., F.N., D.P.O., L.P., M.N.S., S.H., N.V., H.W., R.W., P.W., Z.Z., L.R., J.W., Y.Zh., H.K., U.S., J.S., G.S.; Data were acquired and provided by the IMAGEN and STRATIFY consortiums. Supervision: G.S., J.F., W.L., V.J., S.Pet., C.Z., A.M., J.D., T.J., T.L.. Writing—original draft: Y.X., S.P., G.S. Writing—review & editing: Y.X., G.S., S.P., J.D., S.Pet., L.Q., S.D., Z.Z., J.F., W.L..

## Data availability

The data analysed in this study were obtained from the IMAGEN consortium and associated cohorts. Due to ethical and data protection restrictions, the raw data are not publicly available. Researchers may request access to the data through the official data access procedures of the respective consortia. Processed data supporting the findings of this study are available from the corresponding author upon reasonable request and subject to consortium approval.

## Code availability

The code used to perform the analyses in this study is publicly available at GitHub (https://github.com/Shmily94/Digital_twin_brain-psychiatry-.git). Custom codes developed in PyTorch for personal computing environments and in C++ for high-performance computing (HPC) systems of the DTB platform is available via the DTB Consortium GitHub repository (https://github.com/DTB-consortium/Digital_twin_brain-open) and Zenodo (https://doi.org/10.5281/zenodo.13995756). Please note that parts of the HPC implementation depend on specific hardware and software configurations. Further technical information can be requested from the DTB Consortium via.

## References

1. Dubois, J. & Adolphs, R. Building a Science of Individual Differences from fMRI. Trends Cogn. Sci. 20, 425–443 (2016).

2. Bassett, D. S. & Sporns, O. Network neuroscience. Nat. Neurosci. 20, 353–364 (2017).

3. Sejnowski, T. J., Churchland, P. S. & Movshon, J. A. Putting big data to good use in neuroscience. Nat. Neurosci. 17, 1440–1441 (2014).

4. Huys, Q. J. M., Maia, T. V. & Frank, M. J. Computational psychiatry as a bridge from neuroscience to clinical applications. Nat. Neurosci. 19, 404–413 (2016).

5. Lee, J. H., Liu, Q. & Dadgar-Kiani, E. Solving brain circuit function and dysfunction with computational modeling and optogenetic fMRI. Science 378, 493–499 (2022).

6. Jirsa, V. K., Sporns, O., Breakspear, M., Deco, G. & McIntosh, A. R. Towards The Virtual Brain: network modeling of the intact and the damaged brain. Arch. Ital. Biol. 148, 189–205 (2010).

7. Deco, G. & Kringelbach, M. L. Great expectations: using whole-brain computational connectomics for understanding neuropsychiatric disorders. Neuron 84, 892–905 (2014).

8. Shine, J. M. et al. Computational models link cellular mechanisms of neuromodulation to large-scale neural dynamics. Nat. Neurosci. 24, 765–776 (2021).

9. Lu, W. et al. Imitating and exploring human brain’s resting and task-performing states via resembling brain computing: scaling and architecture. Natl. Sci. Rev. nwae080 (2024) doi:10.1093/nsr/nwae080.

10. Lu, W. et al. Simulation and assimilation of the digital human brain. Nat. Comput. Sci. 4, 890–898 (2024).

11. Braun, U. et al. From Maps to Multi-dimensional Network Mechanisms of Mental Disorders. Neuron 97, 14–31 (2018).

12. Global, regional, and national burden of 12 mental disorders in 204 countries and territories, 1990–2019: a systematic analysis for the Global Burden of Disease Study 2019. Lancet Psychiatry 9, 137–150 (2022).

13. Patel, V. et al. The Lancet Commission on global mental health and sustainable development. The Lancet 392, 1553–1598 (2018).

14. McIntyre, R. S. et al. Treatment-resistant depression: definition, prevalence, detection, management, and investigational interventions. World Psychiatry Off. J. World Psychiatr. Assoc. WPA 22, 394–412 (2023).

15. Kotov, R. et al. The Hierarchical Taxonomy of Psychopathology (HiTOP): A dimensional alternative to traditional nosologies. J. Abnorm. Psychol. 126, 454–477 (2017).

16. Insel, T. R. The NIMH Research Domain Criteria (RDoC) Project: precision medicine for psychiatry. Am. J. Psychiatry 171, 395–397 (2014).

17. Caspi, A. et al. The p Factor: One General Psychopathology Factor in the Structure of Psychiatric Disorders? Clin. Psychol. Sci. J. Assoc. Psychol. Sci. 2, 119–137 (2014).

18. Vanes, L. D. & Dolan, R. J. Transdiagnostic neuroimaging markers of psychiatric risk: A narrative review. NeuroImage Clin. 30, 102634 (2021).

19. Goodkind, M. et al. Identification of a Common Neurobiological Substrate for Mental Illness. JAMA Psychiatry 72, 305–315 (2015).

20. McTeague, L. M. et al. Identification of Common Neural Circuit Disruptions in Cognitive Control Across Psychiatric Disorders. Am. J. Psychiatry 174, 676–685 (2017).

21. Zilverstand, A., Huang, A. S., Alia-Klein, N. & Goldstein, R. Z. Neuroimaging Impaired Response Inhibition and Salience Attribution in Human Drug Addiction: A Systematic Review. Neuron 98, 886–903 (2018).

22. Russo, S. J. & Nestler, E. J. The brain reward circuitry in mood disorders. Nat. Rev. Neurosci. 14, 609–625 (2013).

23. Zanos, P. & Gould, T. D. Mechanisms of Ketamine Action as an Antidepressant. Mol. Psychiatry 23, 801 (2018).

24. Duman, R. S., Sanacora, G. & Krystal, J. H. Altered connectivity in depression: GABA and glutamate neurotransmitter deficits and reversal by novel treatments. Neuron 102, 75–90 (2019).

25. Hu, Y.-T., Tan, Z.-L., Hirjak, D. & Northoff, G. Brain-wide changes in excitation-inhibition balance of major depressive disorder: a systematic review of topographic patterns of GABA- and glutamatergic alterations. Mol. Psychiatry 28, (2023).

26. Ren, Z. et al. Bidirectional Homeostatic Regulation of a Depression-Related Brain State by Gamma-Aminobutyric Acidergic Deficits and Ketamine Treatment. Biol. Psychiatry 80, 457–468 (2016).

27. Biria, M. et al. Cortical glutamate and GABA are related to compulsive behaviour in individuals with obsessive compulsive disorder and healthy controls. Nat. Commun. 14, 3324 (2023).

28. Xie, C. et al. A shared neural basis underlying psychiatric comorbidity. Nat. Med. 29, 1232–1242 (2023).

29. McTeague, L. M., Goodkind, M. S. & Etkin, A. Transdiagnostic impairment of cognitive control in mental illness. J. Psychiatr. Res. 83, 37–46 (2016).

30. Husain, M. & Roiser, J. P. Neuroscience of apathy and anhedonia: a transdiagnostic approach. Nat. Rev. Neurosci. 19, 470–484 (2018).

31. Ritter, C. et al. Evaluation of Prefrontal γ-Aminobutyric Acid and Glutamate Levels in Individuals With Major Depressive Disorder Using Proton Magnetic Resonance Spectroscopy. JAMA Psychiatry 79, 1209–1216 (2022).

32. Marinkovic, K., Alderson Myers, A. B., Arienzo, D., Sereno, M. I. & Mason, G. F. Cortical GABA levels are reduced in young adult binge drinkers: Association with recent alcohol consumption and sex. NeuroImage Clin. 35, 103091 (2022).

33. Prévot, T. & Sibille, E. Altered GABA-mediated information processing and cognitive dysfunctions in depression and other brain disorders. Mol. Psychiatry 26, 151–167 (2021).

34. Zhou, H. et al. Glutamate concentration of medial prefrontal cortex is inversely associated with addictive behaviors: a translational study. Transl. Psychiatry 14, 433 (2024).

35. Godfrey, K. E. M., Gardner, A. C., Kwon, S., Chea, W. & Muthukumaraswamy, S. D. Differences in excitatory and inhibitory neurotransmitter levels between depressed patients and healthy controls: A systematic review and meta-analysis. J. Psychiatr. Res. 105, 33–44 (2018).

36. Sanacora, G., Treccani, G. & Popoli, M. Towards a glutamate hypothesis of depression: An emerging frontier of neuropsychopharmacology for mood disorders. Neuropharmacology 62, 63–77 (2012).

37. Grabski, M. et al. Adjunctive Ketamine With Relapse Prevention–Based Psychological Therapy in the Treatment of Alcohol Use Disorder. Am. J. Psychiatry 179, 152–162 (2022).

38. Dubovsky, S. L. & Marshall, D. Benzodiazepines Remain Important Therapeutic Options in Psychiatric Practice. Psychother. Psychosom. 91, 307–334 (2022).

39. Forsyth, A. et al. Comparison of local spectral modulation, and temporal correlation, of simultaneously recorded EEG/fMRI signals during ketamine and midazolam sedation. Psychopharmacology (Berl.) 235, 3479–3493 (2018).

40. Abdallah, C. G. et al. Ketamine Treatment and Global Brain Connectivity in Major Depression. Neuropsychopharmacology 42, 1210–1219 (2017).

41. Kraguljac, N. V. et al. Ketamine modulates hippocampal neurochemistry and functional connectivity: a combined magnetic resonance spectroscopy and resting-state fMRI study in healthy volunteers. Mol. Psychiatry 22, 562–569 (2017).

42. Breakspear, M. Dynamic models of large-scale brain activity. Nat. Neurosci. 20, 340–352 (2017).

43. Wang, H. et al. Virtual brain twins: from basic neuroscience to clinical use. Natl. Sci. Rev. 11, (2024).

44. Griffiths, J. D., Bastiaens, S. P. & Kaboodvand, N. Whole-Brain Modelling: Past, Present, and Future. in Computational Modelling of the Brain: Modelling Approaches to Cells, Circuits and Networks (eds. Giugliano, M., Negrello, M. & Linaro, D.) 313–355 (Springer International Publishing, Cham, 2022). doi:10.1007/978-3-030-89439-9_13.

45. Hansen, J. Y. et al. Mapping neurotransmitter systems to the structural and functional organization of the human neocortex. Nat. Neurosci. 25, 1569–1581 (2022).

46. Brunel, N. Dynamics of sparsely connected networks of excitatory and inhibitory spiking neurons. J. Comput. Neurosci. 8, 183–208 (2000).

47. Swanson, C. J. et al. Metabotropic glutamate receptors as novel targets for anxiety and stress disorders. Nat. Rev. Drug Discov. 4, 131–144 (2005).

48. Puig, M. V., Watakabe, A., Ushimaru, M., Yamamori, T. & Kawaguchi, Y. Serotonin Modulates Fast-Spiking Interneuron and Synchronous Activity in the Rat Prefrontal Cortex through 5-HT1A and 5-HT2A Receptors. J. Neurosci. 30, 2211–2222 (2010).

49. Celada, P., Puig, M. V. & Artigas, F. Serotonin modulation of cortical neurons and networks. Front. Integr. Neurosci. 7, 25 (2013).

50. William Moreau, A., Amar, M., Le Roux, N., Morel, N. & Fossier, P. Serotoninergic Fine-Tuning of the Excitation–Inhibition Balance in Rat Visual Cortical Networks. Cereb. Cortex 20, 456–467 (2010).

51. Hasselmo, M. E. The role of acetylcholine in learning and memory. Curr. Opin. Neurobiol. 16, 710–715 (2006).

52. Wilson, R. I. & Nicoll, R. A. Endocannabinoid signaling in the brain. Science 296, 678–682 (2002).

53. Kandel, E. R., Koester, J. D., Mack, S. H. & Siegelbaum, S. A. Principles of Neural Science, Sixth *Edition*. (McGraw Hill LLC, 2021).

54. Abbott, L. F. Lapicque’s introduction of the integrate-and-fire model neuron (1907). Brain Res. Bull. 50, 303–304 (1999).

55. Friston, K. J., Mechelli, A., Turner, R. & Price, C. J. Nonlinear responses in fMRI: the Balloon model, Volterra kernels, and other hemodynamics. NeuroImage 12, 466–477 (2000).

56. Isaacson, J. S. & Scanziani, M. How Inhibition Shapes Cortical Activity. Neuron 72, 231–243 (2011).

57. Deco, G. et al. How Local Excitation–Inhibition Ratio Impacts the Whole Brain Dynamics. J. Neurosci. 34, 7886–7898 (2014).

58. Haider, B. & McCormick, D. A. Rapid neocortical dynamics: cellular and network mechanisms. Neuron 62, 171–189 (2009).

59. Yu, Y. et al. A 3D atlas of functional human brain energetic connectome based on neuropil distribution. Cereb. Cortex 33, 3996–4012 (2023).

60. Baroni, F. & Fulcher, B. Synchrony, oscillations, and phase relationships in collective neuronal activity: A highly comparative overview of methods. PLOS Comput. Biol. 21, e1013597 (2025).

61. Suzuki, A., Hara, H. & Kimura, H. Role of the AMPA receptor in antidepressant effects of ketamine and potential of AMPA receptor potentiators as a novel antidepressant. Neuropharmacology 222, 109308 (2023).

62. Dakwar, E. et al. A Single Ketamine Infusion Combined With Motivational Enhancement Therapy for Alcohol Use Disorder: A Randomized Midazolam-Controlled Pilot Trial. Am. J. Psychiatry 177, 125–133 (2020).

63. Riss, J., Cloyd, J., Gates, J. & Collins, S. Benzodiazepines in epilepsy: pharmacology and pharmacokinetics. Acta Neurol. Scand. 118, 69–86 (2008).

64. Haber, S. N. & Knutson, B. The Reward Circuit: Linking Primate Anatomy and Human Imaging. Neuropsychopharmacology 35, 4–26 (2010).

65. Buckner, R. L., Andrews-Hanna, J. R. & Schacter, D. L. The brain’s default network: anatomy, function, and relevance to disease. Ann. N. Y. Acad. Sci. 1124, 1–38 (2008).

66. Cole, M. W. et al. Multi-task connectivity reveals flexible hubs for adaptive task control. Nat. Neurosci. 16, 1348–1355 (2013).

67. Power, J. D. et al. Functional network organization of the human brain. Neuron 72, 665–678 (2011).

68. Lynch, C. J. et al. Frontostriatal salience network expansion in individuals in depression. Nature 633, 624–633 (2024).

69. Reid, A. T. et al. Advancing functional connectivity research from association to causation. Nat. Neurosci. 22, 1751–1760 (2019).

70. Friston, K. J. Functional and Effective Connectivity: A Review. Brain Connect. 1, 13–36 (2011).

71. Jirsa, V. et al. Personalised virtual brain models in epilepsy. Lancet Neurol. 22, 443–454 (2023).

72. Wang, P. et al. Inversion of a large-scale circuit model reveals a cortical hierarchy in the dynamic resting human brain. Sci. Adv. 5, eaat7854 (2019).

73. Stringaris, A. et al. The Brain’s Response to Reward Anticipation and Depression in Adolescence: Dimensionality, Specificity, and Longitudinal Predictions in a Community-Based Sample. Am. J. Psychiatry 172, 1215–1223 (2015).

74. Cole, M. W., Ito, T., Cocuzza, C. & Sanchez-Romero, R. The Functional Relevance of Task-State Functional Connectivity. J. Neurosci. 41, 2684–2702 (2021).

75. Ozeki, H., Finn, I. M., Schaffer, E. S., Miller, K. D. & Ferster, D. Inhibitory stabilization of the cortical network underlies visual surround suppression. Neuron 62, 578–592 (2009).

76. Murphy, B. K. & Miller, K. D. Balanced Amplification: A New Mechanism of Selective Amplification of Neural Activity Patterns. Neuron 61, 635–648 (2009).

77. Joglekar, M. R., Mejias, J. F., Yang, G. R. & Wang, X.-J. Inter-areal Balanced Amplification Enhances Signal Propagation in a Large-Scale Circuit Model of the Primate Cortex. Neuron 98, 222–234.e8 (2018).

78. Murray, J. D. et al. Linking Microcircuit Dysfunction to Cognitive Impairment: Effects of Disinhibition Associated with Schizophrenia in a Cortical Working Memory Model. Cereb. Cortex 24, 859–872 (2014).

79. Yizhar, O. et al. Neocortical excitation/inhibition balance in information processing and social dysfunction. Nature 477, 171–178 (2011).

80. Ahmadian, Y. & Miller, K. D. What is the dynamical regime of cerebral cortex? Neuron 109, 3373–3391 (2021).

81. Bradley, C., Nydam, A. S., Dux, P. E. & Mattingley, J. B. State-dependent effects of neural stimulation on brain function and cognition. Nat. Rev. Neurosci. 23, 459–475 (2022).

## Methods References

1. Mascarell Maričić, L., et al. The IMAGEN study: a decade of imaging genetics in adolescents. Mol. Psychiatry 25, 2648–2671 (2020).

2. Schumann, G. et al. The IMAGEN study: reinforcement-related behaviour in normal brain function and psychopathology. Mol. Psychiatry 15, 1128–1139 (2010).

3. Xie, C. et al. A shared neural basis underlying psychiatric comorbidity. Nat. Med. 29, 1232–1242 (2023).

4. Allen, J. P., Litten, R. Z., Fertig, J. B. & Babor, T. A review of research on the Alcohol Use Disorders Identification Test (AUDIT). Alcohol. Clin. Exp. Res. 21, 613–619 (1997).

5. Kroenke, K., Spitzer, R. L. & Williams, J. B. The PHQ-9: validity of a brief depression severity measure. J. Gen. Intern. Med. 16, 606–613 (2001).

6. Sheehan, D. V. et al. The Mini-International Neuropsychiatric Interview (M.I.N.I.): the development and validation of a structured diagnostic psychiatric interview for DSM-IV and ICD-10. J. Clin. Psychiatry 59 Suppl 20, 22–33;quiz 34-57 (1998).

7. Forsyth, A. et al. Comparison of local spectral modulation, and temporal correlation, of simultaneously recorded EEG/fMRI signals during ketamine and midazolam sedation. Psychopharmacology (Berl.) 235, 3479–3493 (2018).

8. Goodman, R., Ford, T., Richards, H., Gatward, R. & Meltzer, H. The Development and Well-Being Assessment: Description and initial validation of an integrated assessement of child and adolescent psychopathology. J. Child Psychol. Psychiatry 10.1017/S0021963099005909 (2000) doi:10.1017/S0021963099005909.

9. Goodman, R. Psychometric properties of the strengths and difficulties questionnaire. J. Am. Acad. Child Adolesc. Psychiatry 40, 1337–45 (2001).

10. Esteban, O. et al. fMRIPrep: a robust preprocessing pipeline for functional MRI. Nat. Methods 16, 111–116 (2019).

11. Shen, X., Tokoglu, F., Papademetris, X. & Constable, R. T. Groupwise whole-brain parcellation from resting-state fMRI data for network node identification. NeuroImage 82, 403–415 (2013).

12. Shen, X. et al. Using connectome-based predictive modeling to predict individual behavior from brain connectivity. Nat. Protoc. 12, 506–518 (2017).

13. Hansen, J. Y. et al. Mapping neurotransmitter systems to the structural and functional organization of the human neocortex. Nat. Neurosci. 25, 1569–1581 (2022).

14. Lu, W. et al. Imitating and exploring human brain’s resting and task-performing states via resembling brain computing: scaling and architecture. Natl. Sci. Rev. nwae080 (2024) doi:10.1093/nsr/nwae080.

15. Lu, W. et al. Simulation and assimilation of the digital human brain. Nat. Comput. Sci. 4, 890–898 (2024).

16. Brunel, N. Dynamics of sparsely connected networks of excitatory and inhibitory spiking neurons. J. Comput. Neurosci. 8, 183–208 (2000).

17. Binzegger, T., Douglas, R. J. & Martin, K. A. C. A Quantitative Map of the Circuit of Cat Primary Visual Cortex. J. Neurosci. 24, 8441–8453 (2004).

18. Deco, G., Jirsa, V. K. & McIntosh, A. R. Emerging concepts for the dynamical organization of resting-state activity in the brain. Nat. Rev. Neurosci. 12, 43–56 (2011).

19. Isaacson, J. S. & Scanziani, M. How Inhibition Shapes Cortical Activity. Neuron 72, 231–243 (2011).

20. Abbott, L. F. Lapicque’s introduction of the integrate-and-fire model neuron (1907). Brain Res. Bull. 50, 303–304 (1999).

21. Friston, K. J., Mechelli, A., Turner, R. & Price, C. J. Nonlinear responses in fMRI: the Balloon model, Volterra kernels, and other hemodynamics. NeuroImage 12, 466–477 (2000).

22. Yu, Y. et al. A 3D atlas of functional human brain energetic connectome based on neuropil distribution. Cereb. Cortex 33, 3996–4012 (2023).

23. Baroni, F. & Fulcher, B. Synchrony, oscillations, and phase relationships in collective neuronal activity: A highly comparative overview of methods. PLOS Comput. Biol. 21, e1013597 (2025).

24. Deco, G. et al. Resting-state functional connectivity emerges from structurally and dynamically shaped slow linear fluctuations. J. Neurosci. Off. J. Soc. Neurosci. 33, 11239–11252 (2013).

25. Li, J., Kells, P. A., Osgood, A. C., Gautam, S. H. & Shew, W. L. Collapse of complexity of brain and body activity due to excessive inhibition and MeCP2 disruption. Proc. Natl. Acad. Sci. 118, e2106378118 (2021).

